# Seasonal restructuring of heterotrophic microbial communities is differentially affected by glacier type in Greenland fjords

**DOI:** 10.64898/2026.09.22.753439

**Authors:** Marta Mikhno, Lorenz Meire, Renaat Dasseville, Sofie D’hondt, Peter Chaerle, Ilse Daveloose, Koen Sabbe

## Abstract

Accelerated retreat of the Greenland Ice Sheets is increasing freshwater discharge to coastal fjords and promoting transitions from marine-terminating glacier (MTG) regimes to land-terminating (LTG) ones. These changes strongly affect the physical and chemical features of the water column, which in turn impact the structure and functioning of fjord ecosystems. We used high-throughput imaging and DNA metabarcoding approaches to investigate seasonal (spring vs summer) differences in the structure of pelagic microbial communities in two contrasting fjord systems (MTG vs LTG) in southwest Greenland, with focus on the bacterial and heterotrophic protist components, which to date remain understudied in Greenland fjords. Our analyses revealed a marked seasonal restructuring in the microbial communities, especially in the surface layers. In spring, heterotrophic community composition in both fjords manifested a strong link with the diatom spring bloom, being dominated by typical phytoplankton-associated and copiotroph bacterial groups that can utilize algal-derived organic matter, and protists grazing on phytoplankton. In summer, heterotroph communities strongly diverged between the two fjords, most likely caused by the differential impact of MTG vs LTG glaciers on water column structure and chemistry. There was a marked increase in pico- and nano-sized heterotrophs, parasitic protists (dinoflagellate Syndiniales), and bacterial groups indicative of low-nutrient environments. In addition, bacterial, phytoplankton and heterotrophic protist communities become more strongly coupled, either through stronger interactions and/or stronger environmental filtering acting on all groups simultaneously. These observations are indicative of a shift from resource-replete systems in spring to more resourced-depleted systems in summer. However, this shift was less pronounced in the MTG fjord, where subglacial upwelling enabled the persistence of diatom bloom related communities into summer. Finally, we uncovered the presence of a unique, ice mélange associated inner fjord microbial community, which shows similarities with both supraglacial and sea-ice associated communities, in the inner part of the MTG impacted fjord.

## 1 Introduction

Arctic amplification, the accelerated warming of the Arctic relative to the global average, has turned the Arctic into a hotspot of environmental change (Previdi et al., 2021; Rantanen et al., 2022). A major consequence of this warming is the accelerated retreat of the Greenland Ice Sheet (GrIS), which increases glacial freshwater discharge into coastal systems, altering the physical, chemical and biological properties of the water column (Hopwood et al., 2018, 2020).

Fjords, situated at the interface between the ice sheet and the ocean, are among the most productive Arctic ecosystems, acting as important carbon sinks (Rysgaard et al., 2012; Meire et al., 2015) and supporting valuable fisheries (Meire et al., 2017). Their productivity largely depends on the type and magnitude of glacial input. Marine-terminating (or tidewater) glaciers (MTGs) release subglacial meltwater that drives upwelling of nutrient-rich deep waters, leading to sustained phytoplankton blooms throughout summer (Juul-Pedersen et al., 2015; Meire et al., 2017; Hopwood et al., 2018; Cape et al., 2019). In contrast, land-terminating glaciers (LTGs) supply riverine surface runoff that enhances stratification and limits nutrient replenishment, leading to generally lower productivity during summer (Hopwood et al., 2020; Meire et al., 2023). As many tidewater glaciers across Greenland retreat onto land (Joughin et al., 2010; Murray et al., 2015; Catania et al., 2020; Kavan et al., 2025), fjord systems are gradually shifting from MTG- to LTG-dominated regimes. This transition strongly affects fjord hydrography and biogeochemistry which together reshape microbial community composition and functioning (Cameron et al., 2017; Meire et al., 2017, 2023; Wadham et al., 2019; Andresen et al., 2024; Mikhno et al., 2026).

Nuup Kangerlua (NK, Godthåbsfjord) has long been recognized as a sentinel fjord system for studying the impacts of Arctic climate change on coupled cryosphere–marine ecosystem processes (Juul-Pedersen et al., 2015; Meire et al., 2017; Oksman et al., 2022). Extensive research in this fjord, which is dominated by MTGs, has documented how glacial discharge, nutrient fluxes, and fjord circulation respond to ongoing ice-sheet retreat (Mortensen et al., 2011; Meire et al., 2016a; Chua et al., 2025). In contrast, neighbouring Ameralik (AM) is exclusively influenced by runoff from LTGs. The two fjords open into the same coastal water masses, providing an advantageous setting for comparing how glacier type and fjord morphology interact to shape microbial and biogeochemical dynamics. Meire et al. (2023) and Mikhno et al. (2026) showed that both fjords experience seasonal surface freshening during summer due to meltwater input, but temperature regime, nutrient supply, and microbial community structure and productivity diverge. NK maintains colder surface waters and supports higher phytoplankton biomass and primary production throughout summer, largely sustained by the subglacial upwelling of nutrient-rich deep waters. In contrast, AM is characterized by limited nutrient replenishment, and dominance of smaller phytoplankton in warmer, strongly stratified, and turbid surface waters. To date, most studies have focused on the phytoplankton component using traditional microscopy-based approaches (e.g., Krawczyk et al., 2015, 2018; Vonnahme et al., 2025) and recently also (eukaryotic) DNA metabarcoding approaches (Meire et al., 2023; Rodríguez-Marconi et al., 2024; Mikhno et al., 2026). The heterotrophic microbial component of these fjords however remains poorly characterized, particularly the bacterial communities that play pivotal roles in nutrient regeneration, organic matter remineralization, and carbon cycling (e.g., Rokkan Iversen and Seuthe, 2011; Middelboe et al., 2012; Han et al., 2024). Bacteria and heterotrophic protists shape microbial food webs and fluxes, by driving nutrient regeneration (and hence primary production) and energy transfer through the microbial loop (Pomeroy, 1974; Azam et al., 1983). In addition, competition (bacteria) and grazing (heterotrophic protists) also strongly affect phytoplankton dynamics and productivity (Thingstad et al., 1997; Calbet and Landry, 2004). Understanding how heterotrophic microbial communities differ between systems influenced by MTGs and LTGs is therefore crucial for assessing the broader ecological implications of ongoing glacial retreat.

The present study investigates the composition and size structure of pelagic heterotrophic microbial communities in spring and summer in two contrasting fjord systems in southwest Greenland, in order to assess how glacial type and associated environmental conditions shape microbial ecology and ecosystem functioning. Microbial community composition and carbon biomass were characterized using a combination of high-throughput imaging approaches (imaging flow cytometry and FlowCAM) together with 18S and 16S rRNA gene sequencing. More specifically, we tested the following hypotheses: (1) differences in heterotrophic microbial community structure would be less pronounced in spring, when diatom spring blooms occur in both fjords, than in summer, when the differential impact of glacier type on microbial communities and size structure is more prominent; (2) in spring, typical bloom-associated bacterial groups and protist grazers would be dominant, while in summer bacterial communities would shift towards groups associated with post-bloom organic matter processing and protist communities towards smaller pico- and nano-sized organisms under nutrient-depleted conditions; (3) the latter shift would be less pronounced in the MTG-fjord due to continued subglacial upwelling of meltwater, which would enhance nutrient delivery to the phytoplankton and allow for higher phytoplankton growth.

## 2 Material and Methods

### 2.1 Study site and sampling design

Nuup Kangerlua (NK) is a large fjord system in SW Greenland (64°10′N, 51°44′W) (Fig. 1A-D), covering ∼2,013 km² with an average depth of ∼250 m and a maximum depth > 600 m (Mortensen et al., 2011). The main branch extends ∼190 km inland. A major sill (∼170 m depth), located near station GF3 (Fig. 1B), marks the entrance to the fjord (Mortensen et al., 2011). The inner basin receives substantial freshwater, ice, and sediment inputs from three MTGs and three LTGs (Fig. 1B). These inputs generate pronounced spatial and seasonal gradients in salinity, temperature, turbidity, and primary productivity, particularly during spring and late summer, when meltwater discharge and sediment load are at their peak (Mortensen et al., 2013; Krawczyk et al., 2015, 2018; Meire et al., 2016b, 2017).

**Figure 1:**
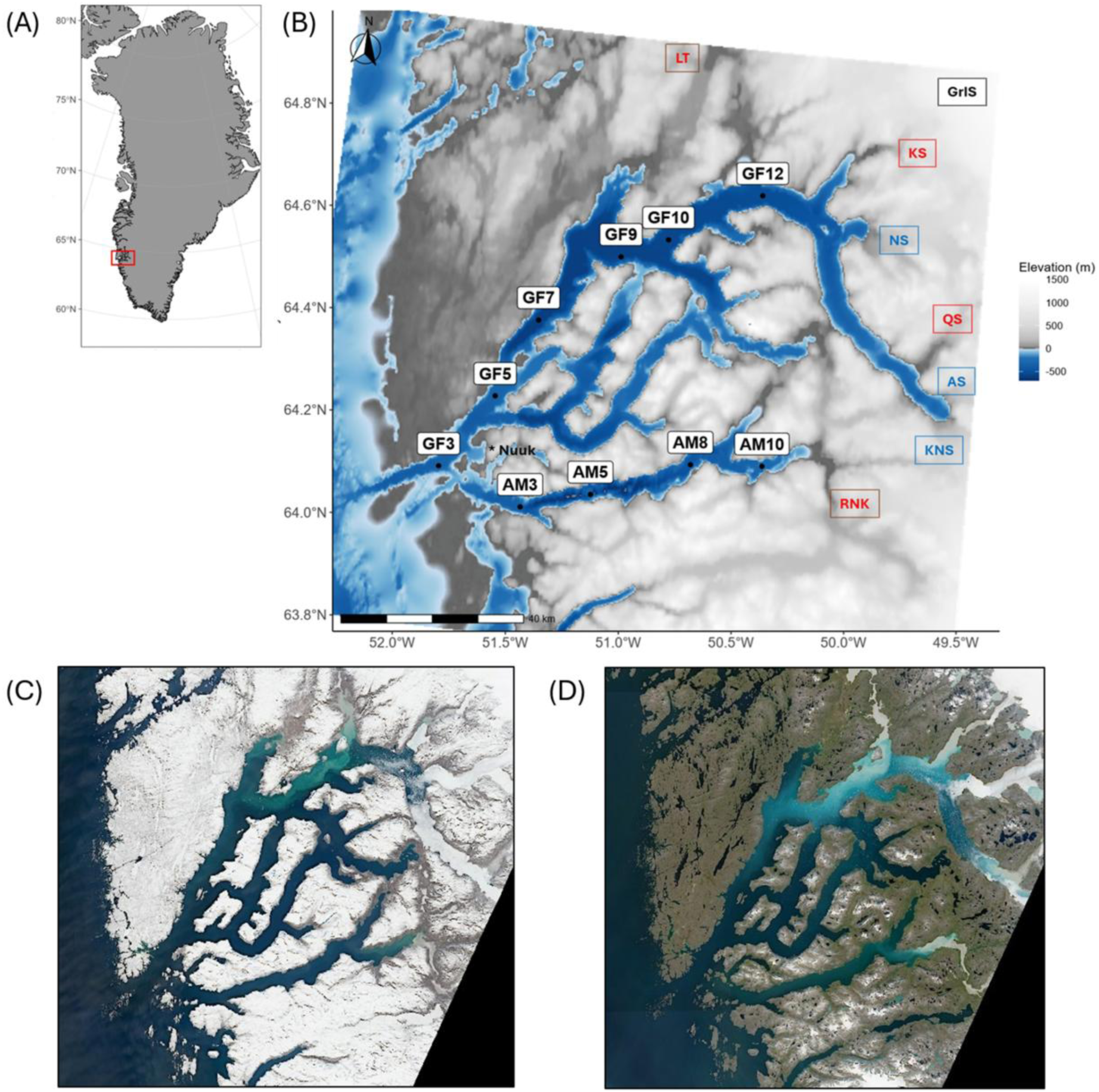
Map of the study area. (A) Geographic context showing the location of the Nuup Kangerlua (NK) and Ameralik (AM) fjords in SW Greenland. (B) Detailed view of the fjords showing sampling stations (circles), with key stations labeled as GFx in NK and AMx in AM. Acronyms indicate freshwater sources: the Greenland Ice Sheet (GrIS); marine-terminating glaciers (MTG, shown in blue: Kangiata Nunaata Sermia [KNS], Akugdlersuup Sermia [AS], and Narsap Sermia [NS]); land-terminating glaciers (LTG, shown in red: Qamanaarssuup Sermia [QS] and Kangilinguata Sermia [KS]); and other freshwater inputs in red, including Lake Tasersuaq (LT) and the LTG-associated River Naajat Kuuat (RNK). (C-D) Sentinel-2 satellite images of the NK and AM fjords in spring (C, 2 June 2022) and summer (D, 31 August 2021). Contains modified Copernicus Sentinel-2 data (ESA), visualized via Sentinel Hub EO Browser (European Space Agency (ESA), 2025).

Ameralik (AM) is located just south of NK, spans ∼400 km², and extends ∼75 km inland (Fig. 1A-D). A sill (∼110 m depth) at the fjord entrance, located just upstream of station AM3 (Fig. 1B), limits water exchange, and the central fjord features deep basins reaching ∼700 m (Stuart-Lee et al., 2021). AM is influenced exclusively by LTGs, with meltwater entering primarily through rivers—most notably the Naajat Kuuat glacial river (RNK)—which generates a marked salinity and turbidity gradient from the inner to the outer basin (Stuart-Lee et al., 2021).

Although located just below the Arctic Circle (∼64°N), the region encompassing NK and AM experiences subzero mean annual air temperatures (−3.9 °C in Nuuk; climate-data.org), and is therefore classified as part of the Arctic domain.

Sampling in both fjords was carried out aboard RV *Avataq* in August 2021 (summer; fig. 1D) and May 2022 (spring; fig. 1C) along longitudinal transects, with 5 summer and 6 spring stations in NK (labelled GFx), and 4 spring and summer stations in AM (labelled AMx). Unless otherwise noted, water was collected using a 5 L Niskin bottle at 1 m (surface); 5 and 10 m (sub-surface); 20 and 40 m (below the deep chlorophyll maximum, bDCM); and 150 and 300 m (deep), as well as at the deep chlorophyll maximum (DCM), determined *in situ* via fluorometric profiles. A table summarizing sampling locations, dates, and DCM depths is provided in the Supplementary Material (Tab. S1).

### 2.2 Environmental data

Vertical profiles of temperature, salinity, density anomaly, turbidity, photosynthetically active radiation (PAR), and chlorophyll fluorescence were acquired using a CTD profiler (SBE 19plus), equipped with a Seapoint Chlorophyll Fluorometer and a Biospherical/Licor Q PAR sensor. Data were collected from the surface down to approximately 5 meters above the seafloor.

For macronutrient analysis, 10 mL subsamples were filtered using 0.45 µm Q-Max GPF syringe filters and immediately frozen at –20°C until processing. Concentrations of nitrate+nitrite (NOx), phosphate (PO₄³⁻), silicate (SiO₂), and ammonium (NH₄⁺) were quantified using standard colorimetric techniques with a Seal QuAAtro autoanalyzer (Graßhoff et al., 2009).

Phytoplankton taxonomic community composition (Mikhno et al., 2026) was integrated into the analyses alongside environmental variables to assess its role in structuring the heterotrophic microbial communities.

### 2.3 Heterotrophic plankton abundance and carbon biomass

The abundance of pico-, nano- and the ≤ 100 µm fraction of microplankton cells was analysed using imaging flow cytometry (iFCM) with an ImageStream®X Mk II. Samples were prepared and analysed as described in Mikhno et al. (2026). Prior to acquisition, samples were stained with SYBR Green I to target nucleic-acid-containing cells. For each event, bright-field imagery and fluorescence signals at 488 nm (SYBR) and 642 nm (chlorophyll autofluorescence) were recorded. Plankton populations were classified in IDEAS® software (v. 6.2) based on their optical and fluorescence properties. In this study, we focused on the heterotrophic component by retaining SYBR-positive cells lacking chlorophyll autofluorescence (i.e., non-autofluorescent cells). These were grouped into size-based functional categories, including bacteria and heterotrophic picoplankton (HP; ≤ 2 µm) as well as small, medium, and large heterotrophic nanoplankton (sHN: 2–3 µm; mHN: 3–5 µm; lHN: 5–20 µm), and heterotrophic microplankton (20–100 µm). Biovolume was estimated following the approach described in Mikhno et al. (2026), the two-dimensional surface area, measured using the IDEAS® imaging software, was multiplied by the cell average width (assuming that width approximates the third spatial dimension, i.e., cell depth; Seelam et al., 2022).

Carbon biomass was subsequently derived from biovolume using established carbon–volume relationships. For the HP fraction, carbon content was estimated using the bacterial conversion proposed by Romanova and Sazhin (Romanova and Sazhin, 2010):

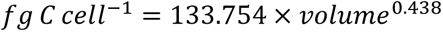

where volume is expressed in µm³. Although the HP fraction may also include heterotrophic picoeukaryotes, a bacterial conversion was applied because the fraction was numerically dominated by bacteria as observed in the iFCM imagery.

For heterotrophic nano- and microplankton groups, carbon content was estimated using the general protist conversion factor of Menden-Deuer and Lessard (Menden-Deuer and Lessard, 2000):

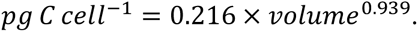

Carbon values were subsequently converted to biomass concentrations and expressed as µg C L⁻¹. Due to technical issues encountered during fieldwork, spring samples from stations AM3 and AM5 were not included in the analysis.

To capture larger heterotrophic particles, seawater (DCM depth only) was additionally concentrated using a 20 µm mesh net and analysed with a FlowCAM® v. 4 imaging system (VisualSpreadsheet® software), which captures the 100–300 µm fraction of the plankton, following the methodology described in Mikhno et al. (2026). Each particle was identified to the lowest possible taxonomic level, and composition and relative abundance were calculated per sample. Images were classified into the following categories: diatoms; the ciliate taxon Tintinnina; the dinoflagellate taxa *Protoperidinium, Tripos* and *Tripos*:part (fragments < 2/3 of the complete cell); zooplankton categories: Copepoda, nauplii, Crustacea and Crustacea:part (fragments of < 2/3 of the organism), Rotifera, Ascidia larvae; non-living material (e.g., detritus, faecal pellets, fibers) was grouped as detrital matter, and unclassified particles were considered as a separate category.

### 2.4 Metabarcoding analysis

For DNA analysis of the pico-, nano- and microplanktonic (< 100 µm) size fractions, 500 mL of seawater was prefiltered through a 100 µm mesh to remove large metazoans and then filtered onto an MF-Millipore MCE membrane filter with a 0.22 µm pore size. Samples were immediately stored at - 80 °C until further processing.

Microbial DNA was extracted from a total of 142 samples (including four negative extraction controls) using the DNeasy PowerLyzer Microbial Kit (Qiagen, Hilden, Germany) following the manufacturer’s protocol. For eukaryotes, the V4 region of the 18S rRNA gene was amplified using the primer set TAReuk454FWD1 (5’-CCAGCASCYGCGGTAATTCC-3’) and the TAReukREV3 (5’-ACTTTCGTTCTTGATYRA-3’) (Stoeck et al., 2010). PCRs and library preparation were done as previously described in Meire et al. (2023). For quality control, blanks and technical triplicate samples were included. Paired-end (2 x 300 base pairs) sequencing was performed with the Illumina MiSeq technology (Illumina, San Diego, US) by Genewiz (Leipzig, Germany). 13,647,731 of raw reads were generated from 150 samples. The high throughput sequence data have been deposited in the NCBI Sequence Read Archive (SRA) under BioProject accession PRJNA1457328.

The 18S Illumina MiSeq data were processed using the DADA2 v. 1.14.1 pipeline (Callahan et al., 2016) in R (R Core Team, 2021) as described in Mikhno et al. (2026). Taxonomic assignment of ASVs was performed using the Protist Ribosomal Reference (PR^2^) database v. 5.0.0 (Vaulot et al., 2023) via the ‘assignTaxonomy’ function, with a minimum bootstrap confidence of 98%. Further ASV processing was conducted using the phyloseq package v. 1.50.0 (McMurdie and Holmes, 2013). Singletons and doubletons were removed to retain genotypes more likely to be ecologically relevant. Metazoa, unidentified ASVs at the kingdom and supergroup levels, and potential contaminants (ASVs with > 1% relative abundance in negative controls) were excluded before downstream analysis. To focus on the heterotrophic microbial community, ASVs belonging to taxonomic classes primarily composed of photosynthetic plankton were excluded from the phyloseq object, while Dinophyceae ASVs were retained on account of the predominance of hetero- and mixotrophic strategies in these organisms, also in Arctic environments (see Mikhno et al., 2026 for more details). The photosynthetic component of the same sample data set was analyzed separately in Mikhno et al. (2026).

For bacteria, the full length 16S rRNA gene was amplified by polymerase chain reaction (PCR) using the primer set: 27F_BCtail-FW (5′-TTTCTGTTGGTGCTGATATTGC_AGAGTTTGATCMTGGCTCAG-3′) and 1492R_BCtail-RV(5′-ACTTGCCTGTCGCTCTATCTTC_CGGTTACCTTGTTACGACTT-3′) (Stackebrandt, 1991). PCRs and library preparation were performed as described in Van Der Loos et al. (2021). For quality control, artificial mock community, blanks and triplicate samples were included. Amplicons for each sample were barcoded using the Oxford Nanopore “PCR Barcoding Expansion Pack 1-96 (EXP-PBC096)”, and subsequently pooled in equimolar ratios and purified using Agencourt AMPure XP beads. The final library was prepared with the ligation-based sequencing kit SQK-LSK109 according to manufacturer’s protocol (Oxford Nanopore Technologies), and sequenced on a MinION with an R9.4.1 flow cell for 72 h. Bases were called with Nanopore’s command line-based tool Guppy, resulting in a total of 15,065,851 reads in 168 samples. Amplicon sequence data have been deposited in the NCBI Sequence Read Archive (SRA) under BioProject accession PRJNA1466576.

The MinION data were visually inspected with NanoPlot (De Coster et al., 2018) and data were demultiplexed with qcat (ONT, https://github.com/nano poretech/qcat). Chimeric reads were removed with Yacrd (Marijon et al., 2020) and the remaining reads were filtered on length (1000–2000 bp) and quality (Q-score *>*10) with NanoFilt (De Coster et al., 2018). The sequences were clustered into Operational Taxonomic Units (OTUs) based on 97% similarity followed by taxonomy assignment via Kraken2 (Lu and Salzberg, 2020). The phyloseq package v. 1.50.0 (McMurdie and Holmes, 2013) was used to further process the OTUs. Singletons and doubletons were removed to focus on genotypes that are more likely to be ecologically relevant. Non-bacterial, chloroplast and mitochondrial OTUs, as well as unidentified OTUs at phylum level and potential contaminants (OTUs with > 1% relative abundance in negative controls) were removed prior to downstream analysis.

### 2.5 Data analysis

All data and statistical analyses, and data visualizations, were performed in R v. 4.4.2 (R Core Team, 2021). Data visualization and plotting was conducted using the ggplot2 package v. 3.5.1 (Wickham, 2016). Relationships between the carbon biomass of plankton size classes and environmental variables were assessed using Spearman rank correlation analyses due to non-normal data distribution (Shapiro– Wilk test). Environmental variables were standardized (*z*-scores) prior to analysis, and pairwise correlations were considered significant at *p* < 0.05. For both heterotrophic protist and bacteria metabarcoding datasets, rarefaction curves were examined via the vegan package (Oksanen et al., 2025). To visualize community composition, bar plots of relative abundances were generated. For protists, ∼99% of reads were represented at the class level, while for bacteria ∼99% was represented at the order level. To assess how community composition was influenced by environmental variables, a constrained analysis of principal coordinates (CAP) was applied to the same datasets using the ‘capscale’ function (Anderson and Willis, 2003) in the vegan package (Oksanen et al., 2025). Bray– Curtis distance matrices were calculated on Hellinger-transformed metabarcoding data, while environmental variables were *z*-score transformed. Environmental parameters considered included Chl a, temperature, salinity, turbidity, PAR, and nutrients. To reduce multicollinearity, variance inflation factors (VIFs) were assessed using the ‘vif.cca’ function, and variables with VIF > 10 were excluded. The final set of constraining variables was selected through forward selection using the ‘ordiR2step’ function from the vegan package (Oksanen et al., 2025). The significance of the CAP model and individual explanatory variables was tested via permutation tests (999 permutations). The top ASVs/OTUs contributing most to the multivariate structure—identified based on the highest loadings on the first two CAP axes—were visualized in ordination plots. Taxa were displayed at the genus level; where classification to this level was not possible, the lowest identified taxonomic level was used. Phytoplankton information was incorporated as passive explanatory variables. A phytoplankton-only phyloseq object was first constructed (see Mikhno et al., 2026), and ASVs were subsequently aggregated into major taxonomic groups as follows: diatoms (Bacillariophyceae, Coscinodiscophyceae, Mediophyceae); green algae (Chlorophyceae, Mamiellophyceae, Pyramimonadophyceae, Trebouxiophyceae); cryptophytes (Cryptophyceae); Phaeocystis (Phaeocystaceae); dictyochophytes (Dictyochophyceae); and chrysophytes (Chrysophyceae). Group-level read abundances were extracted per sample and Hellinger-transformed. These phytoplankton variables were fitted *post hoc* onto the ordination using ‘envfit’ function from the vegan package (Oksanen et al., 2025), allowing the strength and direction of their relationships with heterotrophic community structure to be visualised without affecting the environmentally constrained solution. Only significant (*p* < 0.001, permutation test, n = 999) phytoplankton vectors were displayed on the ordination plot. Environmental variables not retained in the final CAP model were likewise fitted *post hoc* and visualised when significant (*p* < 0.001, permutation test, n = 999). In addition, correlations between dominant bacterial taxa across stations and environmental variables were assessed, and significant relationships (*p* < 0.05) were visualized using the corrplot package (Wei and Simko, 2010). Redundancy analysis (RDA)-based variance partitioning was performed separately for heterotrophic protist and bacterial communities. For heterotrophic protists, the explanatory matrices included environmental variables, phytoplankton communities, and bacterial communities, whereas for bacteria, environmental variables, phytoplankton communities, and heterotrophic protist communities were used as predictors. For this analysis only samples from 1, 5, 10 and DCM depths were included, as depth had a pronounced influence on community composition, and we were mainly interested in the main spatial and seasonal gradients in the surface samples of the data set. Prior to the analyses, multicollinearity among environmental variables was assessed using VIF computed from the RDA models. Variables with VIF values > 10 were excluded to reduce collinearity, and the remaining environmental variables were *z*-score standardised. Variance partitioning was performed using the ‘varpart’ function in the vegan package (Oksanen et al., 2025), based on Hellinger-transformed community data. Separate analyses were conducted for heterotrophic protists and bacteria, and additionally for spring and summer datasets separately. When included as explanatory variables, phytoplankton, bacterial, and heterotrophic communities were first reduced in dimensionality using RDA, and only the first two axes were retained to capture the major gradients in community composition while limiting the number of explanatory variables included in the variance partitioning analyses. The significance of each explanatory block (environment, phytoplankton, and microbial heterotrophs) was assessed using permutation tests (999 permutations) on partial RDA models. To assess co-variation among biotic and abiotic parameters, multiple Mantel tests were performed via the ‘mantel’ function in the vegan package (Oksanen et al., 2025). For this analysis as well, only samples from 1, 5, 10 and DCM depths were included. This method was used to evaluate whether patterns in community composition of the different microbial groups (phytoplankton, heterotrophic protists and bacteria) co-vary with each other and to assess the relationship between each microbial community and environmental factors. A significant Mantel test indicates that as distances between samples increase in one matrix, they also increase in the other, suggesting ecological associations or shared underlying drivers. Co-variability was analysed separately for each fjord and season. First, Bray-Curtis distance matrices of Hellinger-transformed phytoplankton, heterotrophic protist and bacterial data were compared pairwise to assess relationships between microbial communities. Then, each microbial community matrix was compared to Euclidean distance based matrices of *z*-score-transformed environmental parameters. To evaluate the relationship between environmental variables and microbial community composition, a stepwise approach was applied using Mantel tests. First, a matrix including all environmental variables was tested as a single composite predictor (*Environment*). Then, a subset of variables representing nutrients (NOx, PO₄³⁻, SiO₂, and NH₄⁺) was combined and tested as a separate group (*Comb Nutrients*). Finally, each environmental variable was tested individually. The full set of environmental parameters included NOx, SiO₂, PO₄³⁻, NH₄⁺, Chl a, temperature, salinity, turbidity, and PAR. The Mantel test yielded Spearman’s ρ correlation coefficients, and significance was assessed using 999 permutations.

## 3 Results

### 3.1 Spatial and seasonal patterns in heterotrophic microplankton abundance and carbon biomass

#### (i) Microbial heterotrophic plankton < 100 µm (iFCM)

Trends in cell numbers are described in Supplementary Material section S1.

In spring, heterotrophic carbon biomass was highest in inner NK (GF10–GF12), and was largely driven by sHN (Fig. 2A). This fraction averaged ∼14 µg C L⁻¹ in NK compared to ∼9 µg C L⁻¹ in AM and reached its highest values in surface waters (1–5 m) at GF10 (∼40 µg C L⁻¹). HP represented the second most important fraction, with biomass averaging ∼12 µg C L⁻¹ in inner NK (GF10–GF12 average) and ∼8.6 µg C L⁻¹ across the fjord, whereas lower values (∼5 µg C L⁻¹) were observed in inner AM. In contrast, larger heterotrophic fractions contributed only marginally during spring, with mHN averaging ∼2.5 µg C L⁻¹ in both fjords and lHN remaining consistently < 1 µg C L⁻¹. In summer, heterotrophic carbon biomass was substantially higher than in spring, increasing on average from 24 to 212 µg C L⁻¹ (approximately 9-fold), and overall more elevated in AM than in NK, with maxima observed at stations AM5 and AM8 (Fig. 2A). This pattern was again mainly associated with sHN, which remained the dominant heterotrophic fraction in both fjords and averaged ∼150 µg C L⁻¹ in AM compared to ∼100 µg C L⁻¹ in NK. HP represented the second most important fraction, with average biomass of ∼74 µg C L⁻¹ in AM and ∼46 µg C L⁻¹ in NK. A similar fjord contrast was observed for mHN, which averaged ∼40 µg C L⁻¹ in AM and ∼20 µg C L⁻¹ in NK, whereas lHN remained the least abundant fraction, averaging ∼6 µg C L⁻¹ and ∼3 µg C L⁻¹ in AM and NK, respectively. Surprisingly, no heterotrophic microplankton < 100 µm was observed in any of the spring or summer samples.

**Figure 2:**
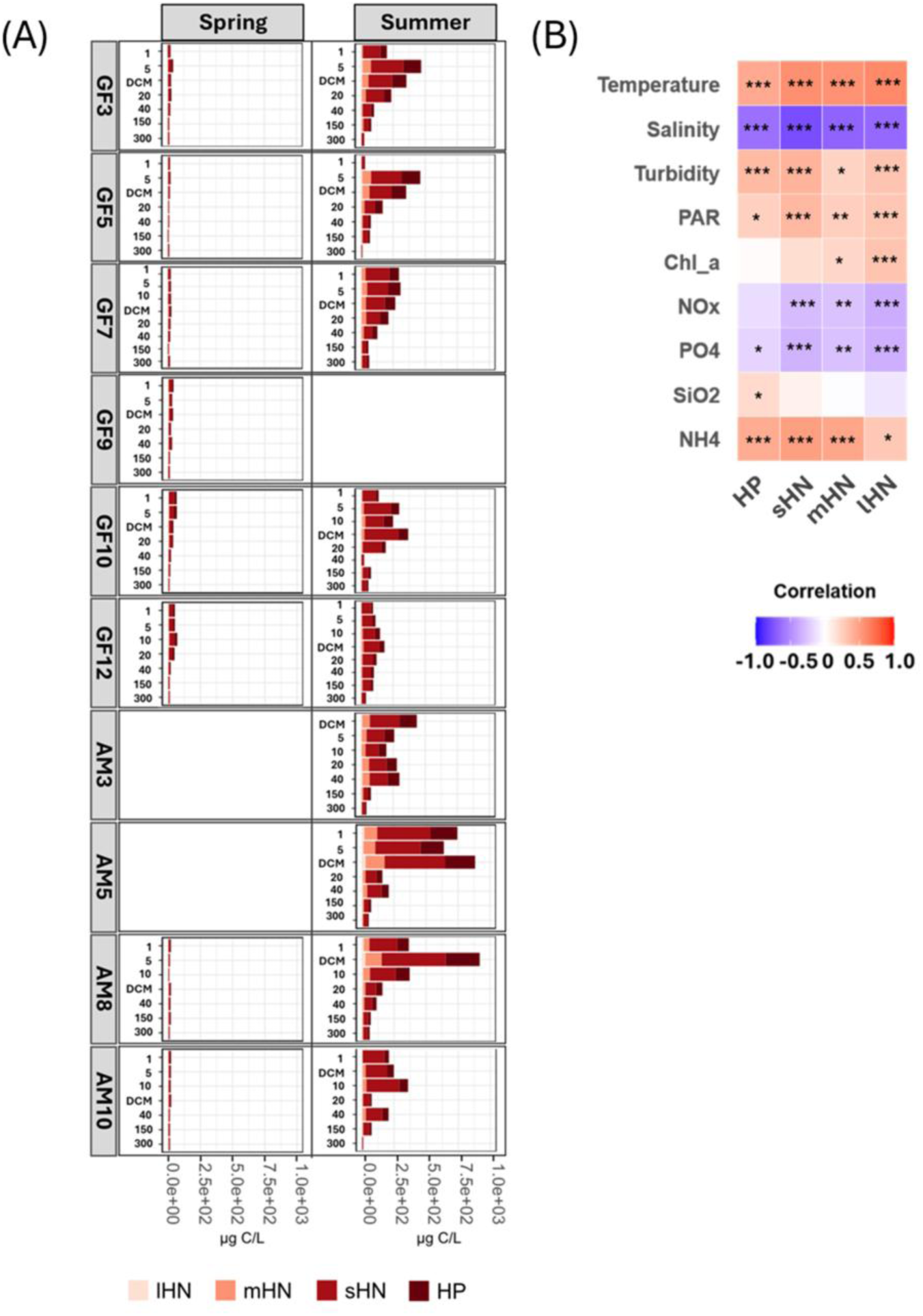
(**A**) Heterotrophic microbial carbon biomass (µg C L^-1^) at discrete sampling depths in NK (GFx) and AM (AMx) in spring (left) and summer (right). DCM = deep chlorophyll maximum. Colors represent microbial plankton functional groups. (**B**) Spearman’s correlation heatmap of microbial plankton functional group carbon biomass and environmental variables. Color intensity indicates strength and direction of correlation (red: positive, blue: negative). Microbial groups: HP – bacteria/picoeukaryotic hetereotrophs, sHN – small nanoheterotrophs, mHN – medium nanoheterotrophs and lHN – large nanoheterotrophs. Significance of permutation tests: ***p ≤ 0.001;**p ≤ 0.005;*p ≤ 0.05.

The carbon biomass of all heterotrophic functional groups was strongly positively correlated with temperature and, to a lesser extent, with turbidity, PAR, and NH₄⁺ (Fig. 2B). Conversely, all groups showed strong negative correlations with salinity and weaker negative correlations with NOx and PO₄³⁻. Only lHN and mHN exhibited significant positive correlations with Chl a.

#### (ii) Microplankton (100-300 µm, FlowCAM)

In spring, the 100–300 µm plankton fraction at the DCM in both fjords was largely dominated by diatoms, whose seasonal composition is described in detail in Mikhno et al. (2026). Tintinnina increased toward the inner AM fjord in spring (from ∼2% at AM3 to ∼9% at AM10) but were rare in NK (< 0.5%, Fig. 3). In summer, Tintinnina reached ∼25% in AM5 but were absent in AM10, while accounting for ∼5% in the NK stations. During summer, *Tripos* and *Protoperidinium* were most abundant at GF5 (6% and 1%, respectively), with *Tripos* also contributing ∼5% at AM10. Copepods reached ∼6.5% at GF5 and GF7, while nauplii were relatively more abundant in AM (∼4% at AM3 and ∼2% at AM5). Crustacea contributed up to ∼4% in NK but were not detected in AM, where only crustacean remains (*crustacea:part*) were observed, reaching ∼9% at AM3. Rotifers were detected at GF5 (∼3%), while in AM they were relatively more abundant, contributing ∼6% at AM3 and ∼5% at AM10. Detrital material dominated the innermost stations of both fjords, exceeding 75% of the total relative particle abundance.

**Figure 3:**
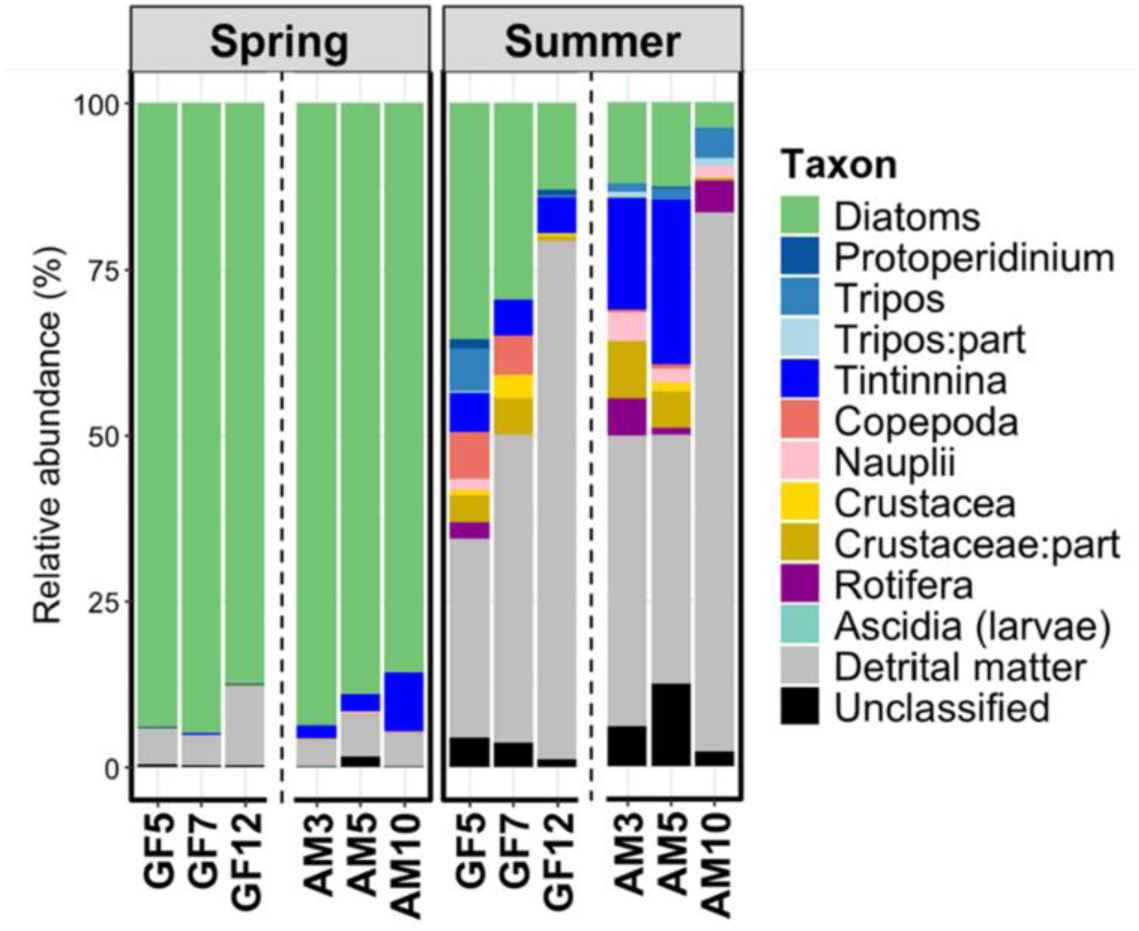
Relative abundances (%) of planktonic and particle groups detected by FlowCAM analysis. The data represent the 100–300 µm plankton fraction from the deep chlorophyll maximum (DCM) during spring (left) and summer (right) in NK (GF5, GF7, GF12) and AM (AM3, AM5, AM10).

### 3.2 Microbial communities: metabarcoding insights and environmental interactions

Inspection of the rarefaction curves for the heterotrophic protist and bacterial metabarcoding data suggests that a significant part of the diversity was captured, as most curves show a declining slope or begin to plateau (Fig. S2a, b). After removal of singletons and doubletons, metabarcoding analysis yielded a total of 3,377 ASVs from 3,613,338 reads across 136 heterotrophic protist samples (with two samples excluded due to technical issues) and 1,942 OTUs from 7,335,115 reads across 138 prokaryotic samples.

#### (i) Heterotrophic protists

Metabarcoding analysis of the heterotrophic protist community revealed pronounced differences between fjords, seasons, and along the depth gradient (Fig. 4). In both seasons, the heterotrophic protist assemblage was strongly dominated by dinoflagellates, primarily represented by the class Dinophyceae (Fig. 4), with the most abundant identified genera included *Gyrodinium* (∼30%), *Heterocapsa* (∼4%), *Gymnodinium* (∼3.5%), *Prorocentrum* (2%) and *Alexandrium* (1.4%; Figs. S3-5). A large fraction of the Dinophyceae reads (∼57%) however could not be assigned to lower taxonomic levels.

**Figure 4:**
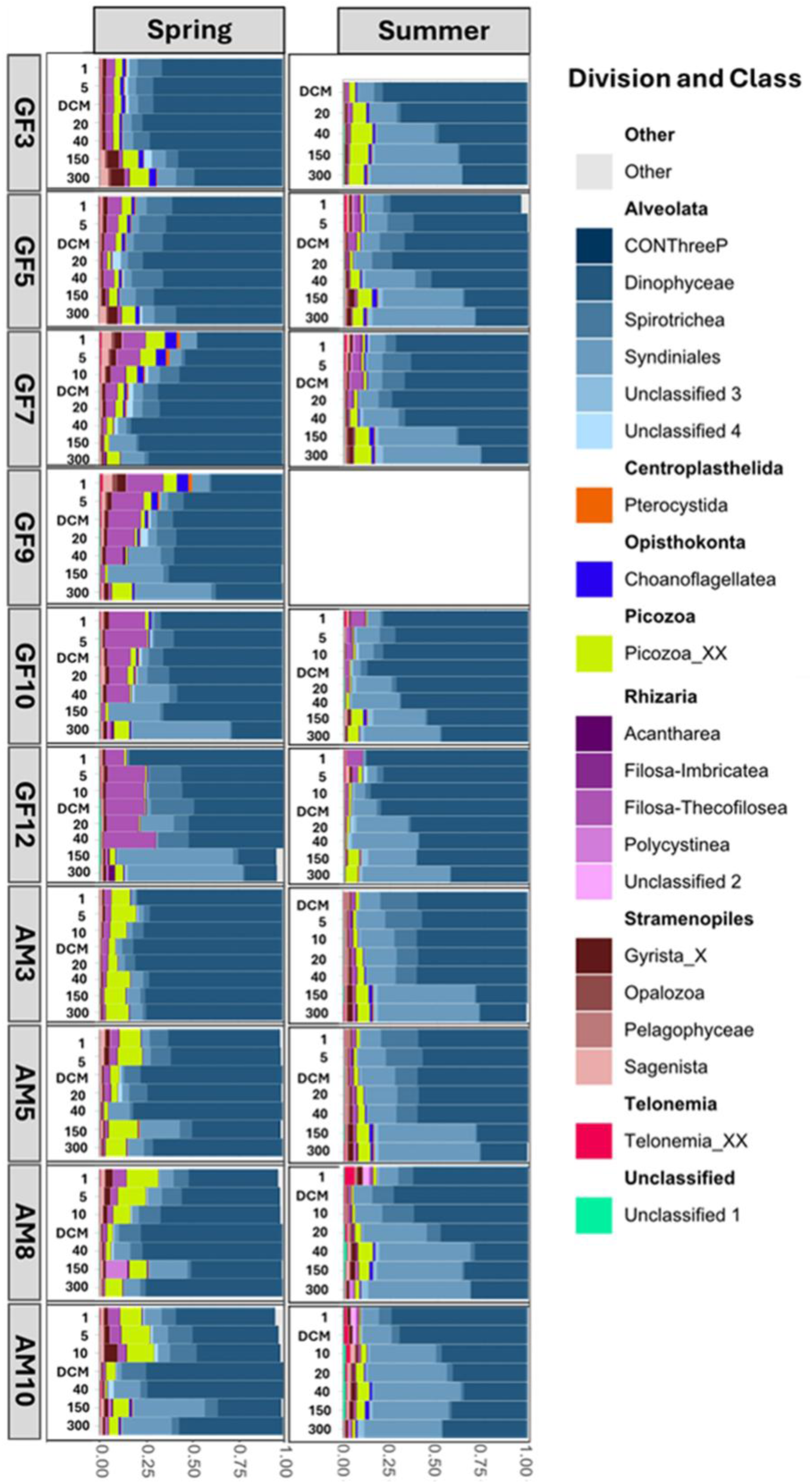
Heterotrophic protist relative abundances by division and 20 most abundant classes (∼99% of total reads) across discrete depths and stations in NK (GFx) and AM (AMx) during spring (left) and summer (right), progressing from outer (GF3, AM3) to inner fjord (GF12, AM10). Other-represent less abundant classes.

In spring, Dinophyceae accounted for an average relative read abundance of ∼60% in NK and ∼65% in AM, with highest values at the fjord mouths, decreasing up-fjord. The main taxa were unclassified Dinophyceae, *Gyrodinium* and unclassified Gymnodiniales (Fig. S4). The class Syndiniales represented the second most abundant dinoflagellate group (∼12% in NK, ∼9% in AM). In both fjords, Syndiniales increased up-fjord and were generally more abundant at greater depths (150–300 m), particularly in the inner fjords. Within the Syndiniales, Dino-Group I displayed a relatively homogeneous distribution throughout the water column in both fjords (except in GF10 and 12 where they were rare), whereas Dino-Group II was largely confined to deeper layers (40–300 m). Other abundant groups included Picozoa, the ciliate class Spirotrichea, and the rhizarian group Filosa–Thecofilosea. Picozoa exhibited higher relative abundances in AM (∼9%) than in NK (∼4%), where they were most abundant in the outer fjord. Spirotrichea showed comparable relative abundances in both fjords (∼8%), with Oligotrichida and Choreotrichida being dominant. Following the PR^2^ taxonomy, which separates Choreotrichida from Choreotrichida*-*Tintinnina, tintinnids were classified separately and represented <1% of relative abundance. Filosa–Thecofilosea, mainly represented by heterotrophic flagellates such as Cryomonadida, were more abundant in NK, where they displayed a clear increase from ∼3% at GF3 to ∼15% at GF12, while accounting for only ∼2.4% in AM.

In summer, Dinophyceae accounted for ∼50% in outer NK and in AM, with their dominance increasing markedly toward inner NK (up to 70% at GF10–GF12). The dominant taxa were unclassified Dinophyceae and *Gyrodinium*, but the dominant ASVs were largely different from those observed in spring. Other abundant genera included *Heterocapsa*, *Gymnodinium*, *Alexandrium*, and, to a lesser extent, *Prorocentrum* (Fig. S5). *Heterocapsa* was notably more abundant in AM than in NK (∼8% vs ∼1%), whereas *Gymnodinium* prevailed in surface waters (1 m) at mid-NK stations (∼12% at GF7– GF10). *Alexandrium* was particularly important at the DCM at GF3 (23%) but was not detected at GF12 (Fig. S5). Syndiniales were relatively more abundant in both fjords compared to spring, particularly below 20 m, averaging 22% in NK and 30% in AM. Dino-Group-I and II dominated the Syndiniales assemblage, with higher clade diversity in summer than in spring (Fig. S5). Picozoa accounted for ∼4% in both fjords. Spirotrichea were relatively more abundant in AM than in NK (∼8% and ∼4%, respectively), peaking at stations AM3-AM8 and being mainly represented by the order Choreotrichida. In NK, their relative abundance was highest in the mid-fjord region, and lowest at GF3 and in the inner fjord. Filosa–Thecofilosea averaged ∼1% of the relative abundance in both fjords. For reference, bar plots illustrating the relative contribution of both phytoplankton and heterotrophic protist groups across stations and depths are provided in fig. S6.

The CAP model (Fig. 5A–B) explained about 44% of the total variation in community composition (adjusted R² = 42%, *p* < 0.001), revealing clear seasonal, spatial and depth patterns. Forward selection retained temperature, salinity, turbidity, PAR, NOx, NH₄⁺ and Chl a as significant explanatory variables. The first axis (CAP1), which explained 16.4% of the variance, was strongly positively correlated with NOx (and PO₄³⁻), and, to a lesser extent, with salinity, and negatively with temperature and to a lesser extent Chl a, turbidity and PAR. This axis mainly captured depth structuring in the communities, separating surface and DCM from deeper waters. Spring surface samples were associated with higher Chl a, especially in NK, whereas summer surface samples aligned with warmer conditions, particularly in AM. Deeper layers were more closely associated with nutrient-rich conditions, and higher salinity. The second axis (CAP2), explaining 15.5% of the variance, was positively correlated with Chl a and to a lesser extent PAR and salinity, and negatively with NH₄⁺, NOx, and temperature. Along this axis, spring samples from colder, phytoplankton (Chl a) rich waters (especially diatoms and *Phaeocystis*, but the latter only in NK, (see Mikhno et al., 2026) were separated from summer samples from warmer waters with higher NH₄⁺ concentrations (especially 20–40 m samples). The distinction between spring and summer samples however decreased with depth. Summer surface samples mainly differed from spring samples by the higher importance of dinoflagellates (*Gymnodinium*, *Heterocapsa*), particularly in AM, and green algae, Cryptophytes, and Dictyochophytes (Fig 5A). The Syndiniales Dino-Group-II-Clade-27 and Dino-Group-II-Clade-10-and-11 and chrysophytes characterized summer samples at 20–40 m in outer NK and inner AM, showing a strong correlation with NH₄⁺. Deep samples (150–300 m) were characterised by unclassified Dinophyceae and *Gyrodinium* in spring, whereas Dino-Group-I-Clade-1 and *Heterocapsa* dominated in summer.

**Figure 5:**
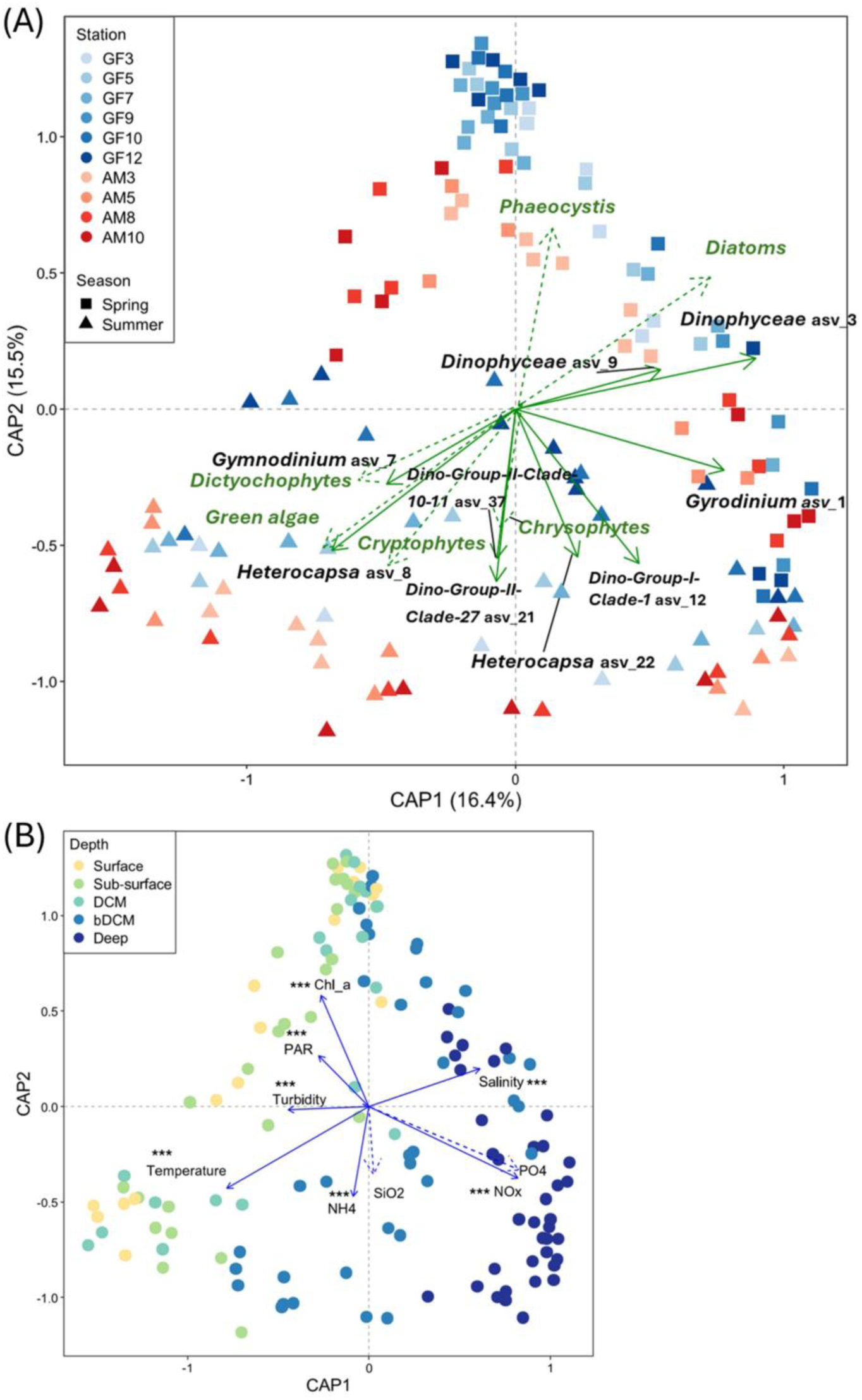
(**A–B**) Constrained analysis of principal coordinates (CAP) ordination of protist communities based on metabarcoding data and environmental variables. (**A**) Samples are colored by station and shaped by season; solid green vectors represent taxa driving multivariate patterns (genus or lowest taxonomic level and ASV number), and dashed green vectors show phytoplankton functional groups passively projected (envfit) onto the ordination. (**B**) Biplot of sample depths: surface (1 m yellow), sub-surface (5, 10 m light-green), deep chlorophyll maximum (DCM, cyan), below DCM (bDCM, 20, 40 m, blue), and deep (150, 300 m, dark blue). Blue vectors indicate environmental gradients, with dashed vectors showing passively fitted variables (envfit). Asterisks denote permutation test significance (*** p ≤ 0.001).

#### (ii) Bacteria

Metabarcoding analysis of prokaryotic communities revealed broadly similar bacterial compositions between the outer and mid sections of NK (GF3–GF9) and the AM stations, with both fjords exhibiting comparable seasonal patterns (Fig. 6; S7-8). In contrast, the inner part of NK (GF10–GF12) harboured a more distinct community, especially in summer. Overall, bacterial communities were consistently dominated by Gammaproteobacteria, Bacteroidia and Alphaproteobacteria, though their relative abundances and specific composition varied spatially and seasonally.

**Figure 6:**
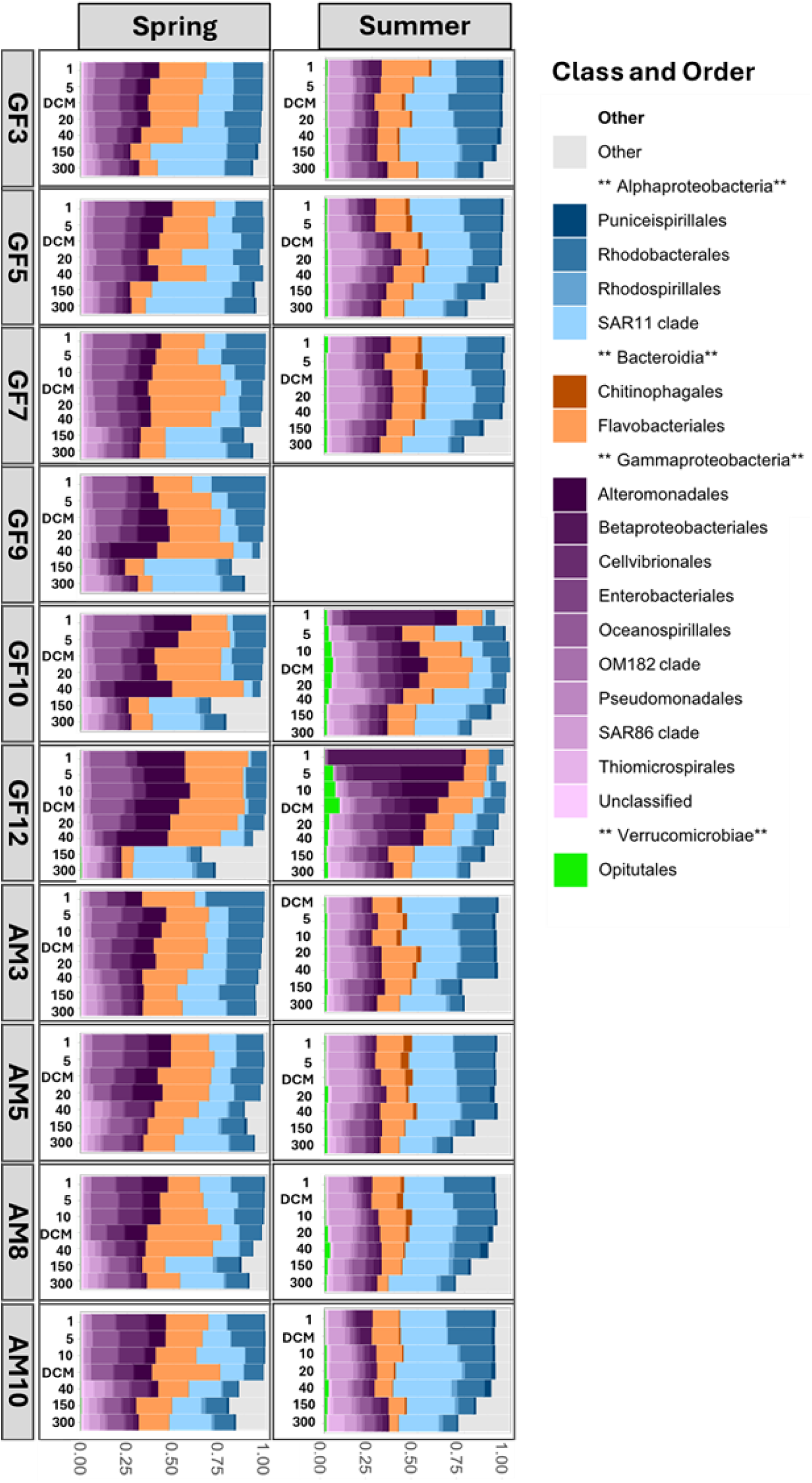
Bacterial relative abundances by class and 17 most abundant orders (∼99% of total reads) across discrete depths and stations in NK (GFx) and AM (AMx) during spring (left) and summer (right), progressing from outer (GF3, AM3) to inner fjord (GF12, AM10). Other- represent less abundant orders.

In spring, Flavobacteriales (Bacteroidia) were the dominant order in the upper 40 m in mid to inner NK and at station AM8, reaching relative abundances of ∼30%. Within this order, *Flavobacterium* was the most abundant genus, followed by *Polaribacter*. In the outer to mid sections of NK and in AM, Gammaproteobacteria accounted for approximately 40% of the bacterial community. The most prevalent orders included Oceanospirales (∼10%), mainly uncultured *Nitrincolaceae*, and *Pseudohongiella* (particularly in deep waters), Cellvibrionales (∼7% in NK, ∼9% in AM), mainly the SAR92 clade, and Alteromonadales (∼7%, predominantly *Colwellia*). *Colwellia* displayed contrasting spatial and depth trends between fjords: in NK, where it was more abundant than in AM, its relative abundance increased up-fjord while in AM it declined, and it decreased markedly below 40 m. Other taxa such as Pseudomonadales, the SAR86 clade, and Betaproteobacteriales were present in lower abundances (<3%) and showed similar distributions in both fjords, with SAR86 clade and OM43 clade (Betaproteobacteriales) being more important in deeper waters. In the inner stations of NK (GF10– GF12), Gammaproteobacteria increased to ∼45%, with Alteromonadales as the dominant order (∼15%), followed by Oceanospirales (∼13%) and Cellvibrionales (∼5%), all represented by the same families and genera as in the rest of the fjord. Alphaproteobacteria were largely dominated by the SAR11 Clade Ia, which was abundant in the upper 40 m in outer NK (GF3–GF5) and AM (∼15%) but declining markedly towards inner NK (∼3% in GF12). However, the clade remained dominant at depth (150–300 m), reaching ∼35% in outer NK and ∼20% in other stations. At these depths, SAR11 Clade II also became more prominent, reaching ∼5% in both fjords. Rhodobacterales (Alphaproteobacteria) were more evenly distributed across both fjords, contributing up to ∼14% (except in GF12, ∼8%). This order was primarily composed of the genera *Sulfitobacter* and *Planktomarina*. Deeper waters in the inner fjord regions of both systems were more distinct, with increased contributions (1–5%) from less dominant groups such as Deltaproteobacteria, Dehalococcoidia, and Acidimicrobiia.

In summer, bacterial community composition was quite similar in the outer to mid sections of NK (stations GF3–GF7) and AM, being dominated by Alphaproteobacteria (∼45%), especially SAR11 Clade Ia (∼ 20%). Inner AM was characterized by higher importance of SAR11 Clade IV (∼10% at AM10), which declined down-fjord. In contrast, SAR11 Clade IV was less prevalent in NK (∼2%, mainly outer to mid-fjord). Rhodobacteriales, particularly *Planktomarina* and *Sulfitobacter*, were consistently present and relatively abundant across all stations, each contributing ∼ 5%. Gammaproteobacteria were represented by the same key orders observed in spring, though in lower abundances. Oceanospirales (∼5%)—mainly uncultured *Nitrincolaceae* and *Pseudohongiella*—and Cellvibrionales (∼3%)— mainly SAR92 clade—remained prominent. In contrast, Alteromonadales decreased markedly (∼1%). Additionally, the SAR86 clade increased in summer, reaching ∼10%. Pseudomonadales also showed an increase in summer, reaching ∼2% in both fjords (*Pseudomonas*). Bacteroidia showed a modest decline (to ∼15%) compared to spring. Flavobacteriales remained the dominant order within this group (∼13%), with *Flavobacterium* and *Polaribacter* as the most common genera, with the latter particularly abundant in the inner NK but absent in AM. Chitinophagales were more abundant in summer (∼2%), with uncultured Chitinophagales particularly abundant in the outer region of both fjords but decreasing up-fjord. Alongside changes in dominant classes, Verrucomicrobiae also increased in summer, particularly the order Opitutales, with *Lentimonas* emerging as a notable genus in inner NK but being completely absent in AM.

The inner part of NK—particularly the surface layer—was markedly different from all other stations in both fjords (Fig. 6). At 1 m depth in GF12, Gammaproteobacteria dominated, with *Burkholderiaceae* (order Betaproteobacteriales) reaching up to 70%, esepcially *Polaromonas* (∼20%) and the RS62 marine group (∼7%). Alteromonadales formed the second most dominant order in this area, with *Glaciecola* accounting for ∼20% in GF10 and GF12. Other genera such as *Paraglaciecola* and *Colwellia*, although present across the fjord, were particularly enriched in GF10 and GF12 and almost absent in AM. In the innermost part of NK, however, SAR11 Clade Ia was nearly absent at the surface (<1% at 1 m) but increased significantly with depth, reaching 26% at 40 m.

The CAP model (Fig. 7A–B) explained 56% of the total variation in bacterial community composition (adjusted R² = 55%, *p* < 0.001). Environmental variables selected through forward selection included temperature, salinity, NOx and NH₄⁺. The first axis accounted for 31.4% of the variance, and was strongly negatively correlated with NOx (and PO₄³⁻), and to a lesser extent NH₄⁺ and temperature, and positively with Chl a (mainly diatoms, especially in mid-inner NK and AM8 at 20–40 m depth, and in NK only also *Phaeocystis*). This axis mainly separated the spring surface samples from the deep samples (150-300 m) and to a lesser degree the summer surface samples, which are also segregated from the spring surface samples along the second axis. The summer surface samples from GF10 and GF12 deviate from the other summer samples, while the summer sub-surface samples (5-10 m) from these stations showed a closer association with the spring cluster. The second axis (15.8% of the variation) was strongly positively associated with salinity and to a lesser extent with NOx (and PO₄³⁻), and strongly negatively with temperature. This axis reflects a depth gradient, from warmer, less saline, surface waters to colder, more saline and nutrient-rich deep waters, but also a distinct spring-summer contrast between the surface layers of the two fjords. Spring samples from 1–40 m depth in outer-mid NK and in AM were characterized by uncultured *Nitrincolaceae*, *Colwellia*, *Paraglaciecola,* the SAR92 clade, *Flavobacterium* and *Polaribacter*. Inner NK samples were characterized by elevated abundances of especially *Nitrincolaceae, Colwellia* and *Paraglaciecola*. Summer samples from 1–40 m depth were characterized by the SAR86 clade and SAR11 Clade IV, with both groups being relatively more abundant in AM than NK. Chrysophytes, cryptophytes and green algae were correlated with this summer cluster, while surface samples from inner NK especially correlated with dictyochophytes. Deep-water samples (150–300 m) formed a distinct, stable cluster that did not exhibit clear separation by fjord or season. They were characterized by SAR11 Clade II, the SUP05 cluster, and the NS9 marine group. SAR11 Clade Ia was important in both summer and deep-water communities.

**Figure 7:**
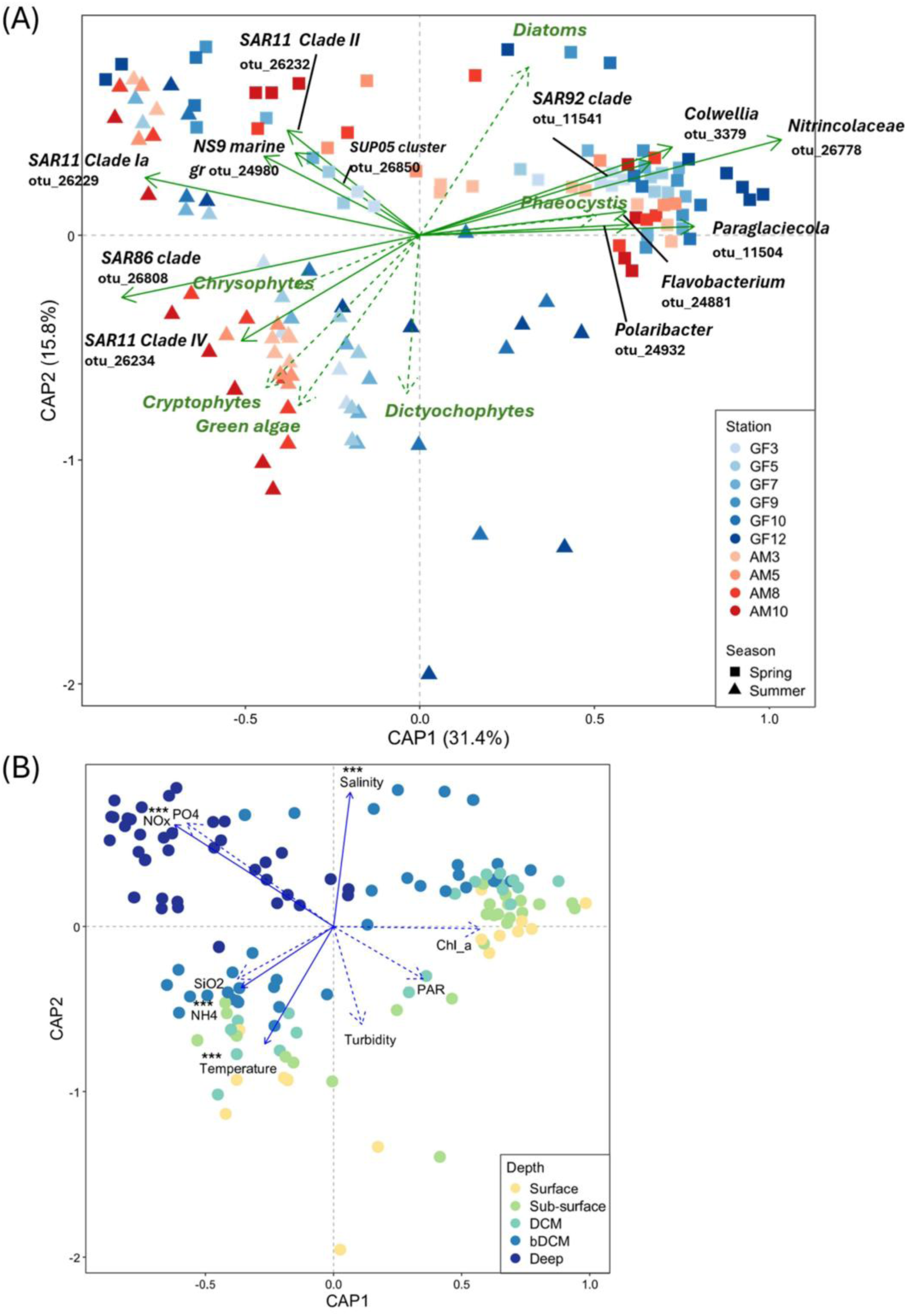
(**A-B**) Constrained analysis of principal coordinates (CAP) ordination showing bacteria community variation based on metabarcoding data and environmental variables. (**A**) Samples are colored by station and shaped by season; solid green vectors represent taxa driving multivariate patterns (genus or lowest taxonomic level and OTU number), and dashed green vectors show phytoplankton functional groups passively projected (envfit) onto the ordination. (**B**) Biplot of sample depths: surface (1 m yellow), sub-surface (5, 10 m light-green), deep chlorophyll maximum (DCM, cyan), below DCM (bDCM, 20, 40 m, blue), and deep (150, 300 m, dark blue). Blue vectors indicate environmental gradients, with dashed vectors showing passively fitted variables (envfit). Asterisks denote permutation test significance (*** p ≤ 0.001).

A Pearson’s correlation analysis illustrating the relationships between the most abundant bacterial taxa and environmental variables is shown in Figure S9. *Flavobacterium* and *Polaribacter* were the only genera positively correlated with Chl a. These two taxa, along with members of *Nitrincolaceae*, *Colwellia*, and the SAR92 clade, also showed negative correlations with temperature. In contrast, *Glaciecola*, *Lentimonas*, *Polaromonas*, and the RS62 marine group were negatively correlated with salinity but positively correlated with turbidity and silicate. SAR Clade IV, *Planktomarina*, NS5 marine group, and *Saprospiraceae* exhibited strong positive correlations with temperature, negative correlations with salinity, and positive correlations with NOx and turbidity. Similarly, SAR Clade Ia was positively correlated with temperature and inorganic nutrients (NH₄⁺, NOx, PO₄³⁻), while negatively correlated with Chl a and PAR. The SUP05 cluster, SAR Clade II, *Pseudohongiella*, and NS9 marine group were strongly positively correlated with NOx and PO₄³⁻, and negatively correlated with Chl a and PAR.

### 3.3 Relationships between biotic (bacteria, heterotrophic and phototrophic protists) and abiotic variables

Environmental variables included in the variance partitioning analyses were temperature, salinity, turbidity, PAR, NOx, NH₄⁺, and Chl a. All variance partitioning models were statistically significant (permutation tests, *p* = 0.001). Bacterial community composition showed a higher overall proportion of explained variance than heterotrophic protists, particularly in summer (Fig. 8A-D). Across both seasons, environmental factors accounted for the largest unique fraction of explained variance for both heterotrophic protists and bacteria, while the unique contributions of the other microbial groups (phytoplankton and respectively bacteria and heterotrophic protists) were consistently smaller. For heterotrophic protists, the total proportion of explained variance was similar between spring and summer (Fig. 8A–B). In spring, a substantial fraction of variance was jointly explained by bacteria and phytoplankton (20%), followed by the shared contribution of environment and bacteria (10%). In summer, the largest fraction of heterotrophic protist variance was jointly explained by all three groups of variables (19%). In spring, a large fraction of bacterial variance was jointly explained by phytoplankton and heterotrophic protists (20%), followed by the shared contribution of environment and heterotrophic protists (13%). Remarkably, in summer, more than half of the explained variance was shared among all three groups (54%), followed by environment and phytoplankton shared contribution (9%).

**Figure 8.**
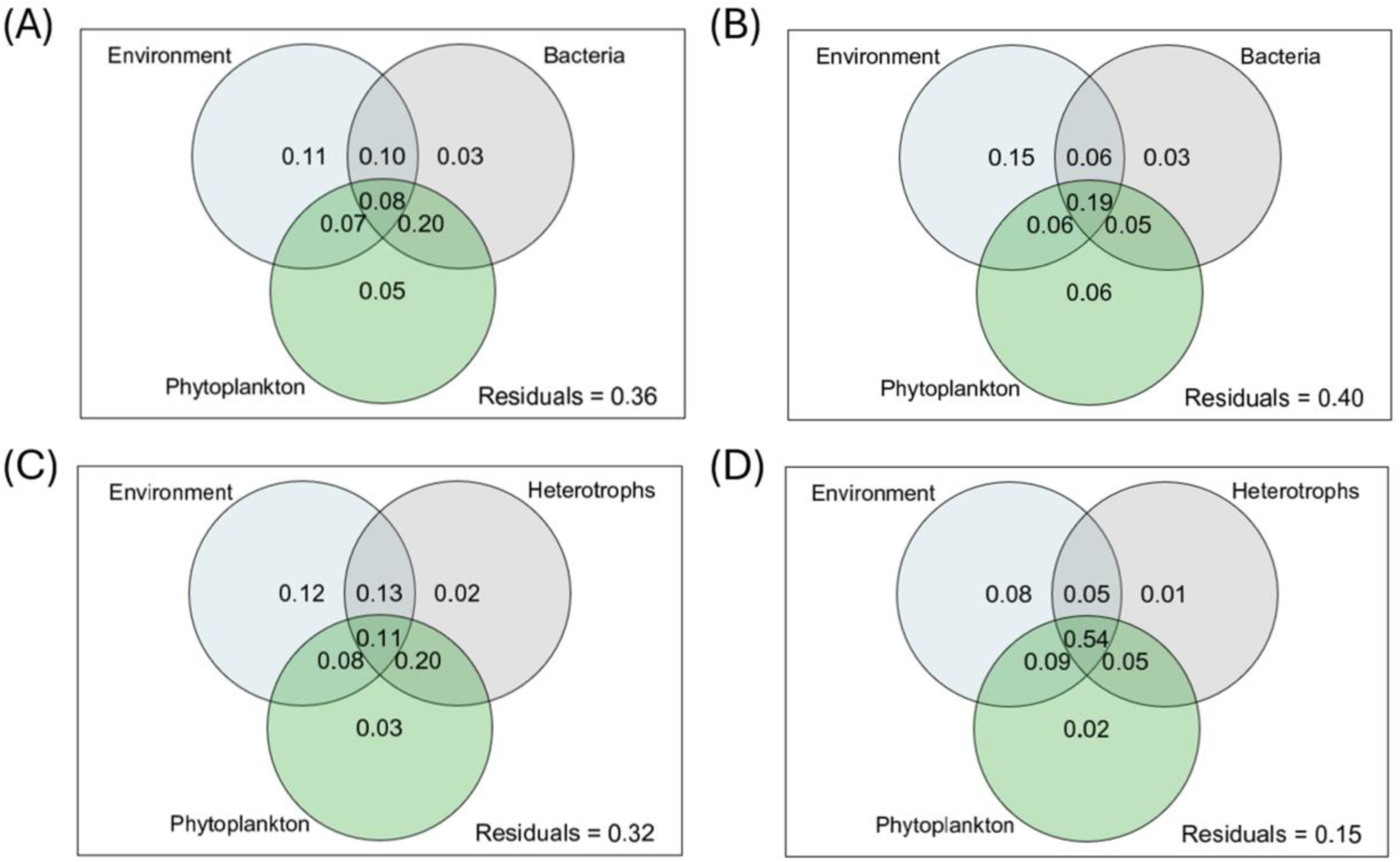
(A–D) Variance partitioning of surface (1, 5, 10 m and DCM depths) community composition (ASVs and OTUs) showing the relative contributions of environmental variables, phytoplankton, and microbial compartments (heterotrophic protists and bacteria). Panels show results for (**A**) heterotrophic protists in spring, (**B**) heterotrophic protists in summer, (**C**) bacteria in spring, and (**D**) bacteria in summer. Values indicate the proportion of explained variance attributed to each unique and shared fraction; unexplained variance is reported as residuals. All models were statistically significant (permutation tests, p = 0.001).

Mantel tests showed that, in both fjords and seasons, surface (1 m–DCM) biological communities were more strongly correlated with each other than with environmental variables (Tab. 1), with inter-community correlations being generally more pronounced in summer. Likewise, correlations between microbial communities and environmental parameters were stronger in summer than in spring across both fjords, with the highest values observed in NK. In NK during spring, salinity emerged as the dominant environmental correlate (ρ > 0.6) across microbial compartments, while in summer, turbidity showed the strongest correlations, particularly with bacteria (ρ > 0.8). Salinity, silicate and PAR were also strongly correlated with heterotrophic protists and bacteria, while its association with phytoplankton was weaker. In AM during spring, phytoplankton and heterotrophic protist communities but not bacteria were strongly correlated with temperature and salinity (ρ > 0.6). As in NK, environmental coupling intensified in summer, when nutrients—particularly silicate—as well as salinity, turbidity, and temperature were strongly correlated with both protist and bacterial communities. Phytoplankton exhibited more moderate correlations with the same environmental variables, but also with NOx and PO₄³⁻.

**Table 1:**
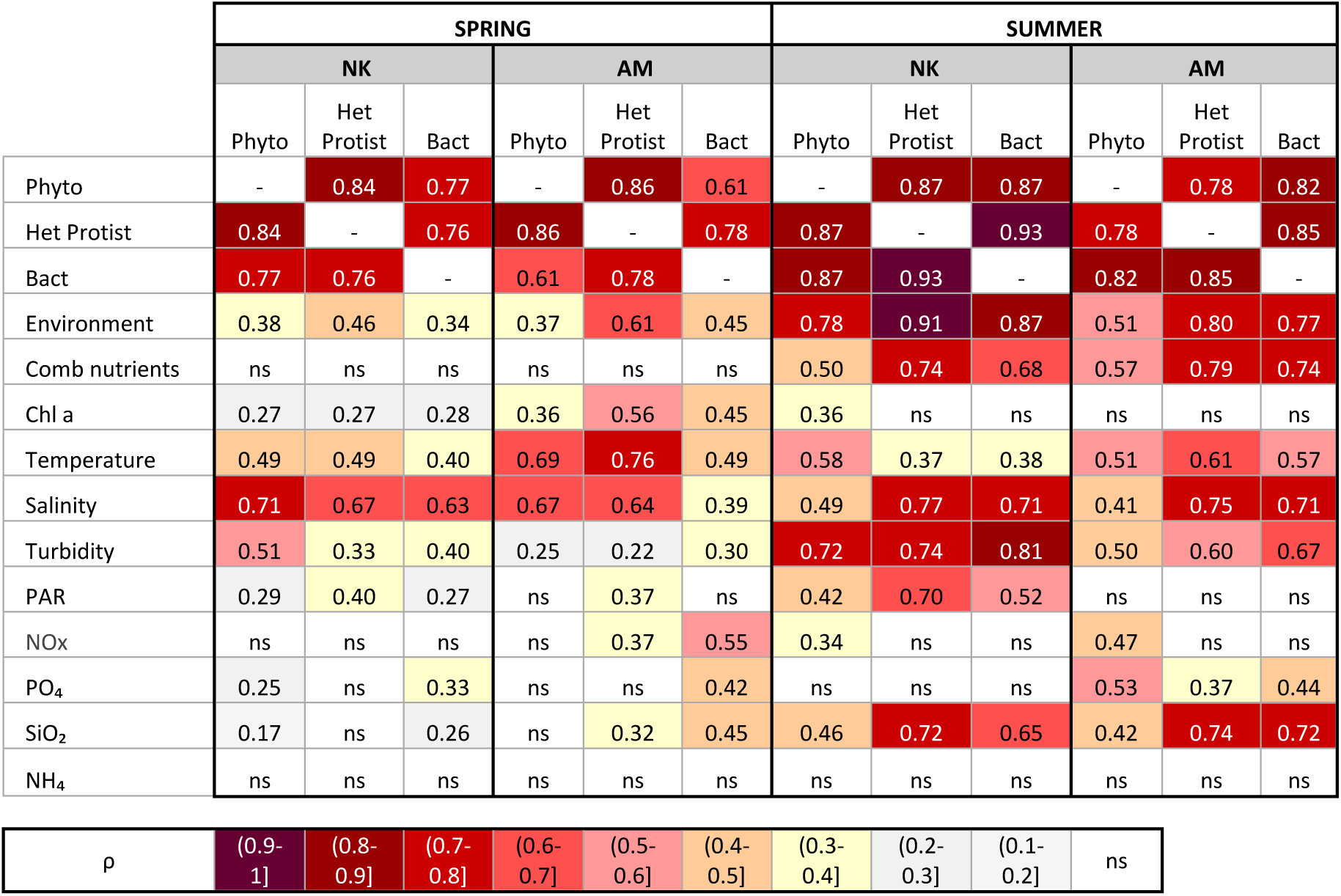
Mantel test correlation coefficients (ρ) for pairwise comparisons among Phyto (phytoplankton), Het protist (heterotrophic protists), and Bact (bacteria), and between each microbial community and environmental variables at selected depths (1, 5, 10 m, and the DCM). Analyses were performed separately for each fjord and season. Significant correlations indicate significant co-variation between the compared datasets. p ≤ 0.05; ns, not significant

|  | SPRING |  |  |  |  |  | SUMMER |  |  |  |  |  |
| --- | --- | --- | --- | --- | --- | --- | --- | --- | --- | --- | --- | --- |
|  | NK |  |  | AM |  |  | NK |  |  | AM |  |  |
|  | Phyto | Het Protist | Bact | Phyto | Het Protist | Bact | Phyto | Het Protist | Bact | Phyto | Het Protist | Bact |
| Phyto | - | 0.84 | 0.77 | - | 0.86 | 0.61 | - | 0.87 | 0.87 | - | 0.78 | 0.82 |
| Het Protist | 0.84 | - | 0.76 | 0.86 | - | 0.78 | 0.87 | - | 0.93 | 0.78 | - | 0.85 |
| Bact | 0.77 | 0.76 | - | 0.61 | 0.78 | - | 0.87 | 0.93 | - | 0.82 | 0.85 | - |
| Environment | 0.38 | 0.46 | 0.34 | 0.37 | 0.61 | 0.45 | 0.78 | 0.91 | 0.87 | 0.51 | 0.80 | 0.77 |
| Comb nutrients | ns | ns | ns | ns | ns | ns | 0.50 | 0.74 | 0.68 | 0.57 | 0.79 | 0.74 |
| Chl a | 0.27 | 0.27 | 0.28 | 0.36 | 0.56 | 0.45 | 0.36 | ns | ns | ns | ns | ns |
| Temperature | 0.49 | 0.49 | 0.40 | 0.69 | 0.76 | 0.49 | 0.58 | 0.37 | 0.38 | 0.51 | 0.61 | 0.57 |
| Salinity | 0.71 | 0.67 | 0.63 | 0.67 | 0.64 | 0.39 | 0.49 | 0.77 | 0.71 | 0.41 | 0.75 | 0.71 |
| Turbidity | 0.51 | 0.33 | 0.40 | 0.25 | 0.22 | 0.30 | 0.72 | 0.74 | 0.81 | 0.50 | 0.60 | 0.67 |
| PAR | 0.29 | 0.40 | 0.27 | ns | 0.37 | ns | 0.42 | 0.70 | 0.52 | ns | ns | ns |
| NO<sub>x</sub> | ns | ns | ns | ns | 0.37 | 0.55 | 0.34 | ns | ns | 0.47 | ns | ns |
| PO<sub>4</sub> | 0.25 | ns | 0.33 | ns | ns | 0.42 | ns | ns | ns | 0.53 | 0.37 | 0.44 |
| SiO<sub>2</sub> | 0.17 | ns | 0.26 | ns | 0.32 | 0.45 | 0.46 | 0.72 | 0.65 | 0.42 | 0.74 | 0.72 |
| NH<sub>4</sub> | ns | ns | ns | ns | ns | ns | ns | ns | ns | ns | ns | ns |

|  |  |  |  |  |  |  |  |  |  |  |
| --- | --- | --- | --- | --- | --- | --- | --- | --- | --- | --- |
| $\rho$ | (0.9-1] | (0.8-0.9] | (0.7-0.8] | (0.6-0.7] | (0.5-0.6] | (0.4-0.5] | (0.3-0.4] | (0.2-0.3] | (0.1-0.2] | ns |

## 4 Discussion

Using a combination of high-throughput imaging and DNA metabarcoding, we performed a detailed seasonal (spring vs summer) comparison of the heterotrophic component of pelagic microbial communities in two adjacent Arctic fjords under the influence of different glacier types (predominantly MTG vs LTG). While (1) we observed a general dominance of Gamma- and Alphaproteobacteria, Bacteroidia and dinoflagellates across seasons and depths in both fjord systems, there was (2) a marked seasonal restructuring in heterotrophic community composition at lower taxonomic levels, which was especially pronounced in the surface layers (1-40m), and which coincided with an on average 9-fold increase in the C biomass of pico- and nano-sized heterotrophs, especially in the LTG fjord. While spring was mainly dominated by Gymnodiniales and copiotroph bacterial groups (e.g. Flavobacteriales), summer saw a notable increase in the parasitic dinoflagellate group Syndiniales, and oligotrophic bacterial groups [e.g. SAR86 (Gammaproteobacteria) and SAR11 clade IV (Alphaproteobacteria)]. (3) The seasonal restructuring augmented the differences in community structure between the MTG and LTG fjord, with the LTG fjord shifting towards a more recycling-dominated microbial food web, whereas the MTG fjord maintained features associated with sustained diatom production. (4) These summertime differences were most pronounced in the inner fjord stations, which were most directly exposed to glacier (MTG) or glacial river (LTG) impact, with unique, previously unreported sea-ice like communities in the inner MTG fjord. (5) In both fjords, there was a clear depth structuring in the communities, which became more pronounced in summer as stratification stabilized the vertical hydrographic structure of the water column, except in mid to inner NK, where subglacial upwelling persisted. (6) Finally, variation partitioning and Mantel tests revealed a marked increase in environmental filtering and/or biotic coupling between bacterial, phytoplankton and heterotrophic protist communities in summer, likely related to a shift from resource-replete spring to resource-depleted summer conditions. Below, these findings are discussed in more detail.

### 4.1 Phytoplankton blooms shape spring heterotrophic microbial community composition

The occurrence of extensive diatom-dominated blooms in both fjords (Mikhno et al., 2026) was reflected in overall rather similar bacterial and heterotrophic protist communities in the fjords, mainly dominated by dinoflagellates (Dinophyceae), Gammaproteobacteria, Bacteroidia, and Alphaproteobacteria. However, there were also differences between the two fjords, with Filosa-Thecofilosea flagellates being important in NK, and Picozoa being more characteristic of AM. This could be related to subtle differences in phytoplankton community structure between the fjords (presence of haptophyte *Phaeocystis* in NK vs green algae, Crypto- and Chrysophyceae in AM).

In spring, dinoflagellate communities were mainly composed of the class Dinophyceae, especially the genus *Gyrodinium* from the athecate order Gymnodiniales. This is in agreement with previous reports of Gymnodiniales dominance in both fjords in spring, based on microscopy (Engel Arendt et al., 2010; Krawczyk et al., 2018) and DNA metabarcoding (Meire et al., 2023). More exact identification of Gymnodiniales was not possible, due to the limited phylogenetic resolution of the 18S rDNA marker, especially for distinguishing closely related gymnodinioid lineages (Saldarriaga et al., 2001), but also because of the general lack of reference sequences. It also has to be kept in mind that dominance of dinoflagellates in metabarcoding data sets should be interpreted with some caution, as dinoflagellates are known to possess high rRNA gene copy numbers, which can lead to their overrepresentation in amplicon-based datasets (Lin, 2006; Ruvindy et al., 2023). In NK, previous metabarcoding-based observations of dinoflagellate dominance contrasted with substantially lower abundances inferred from microscopy (Rodríguez-Marconi et al., 2024). Ciliates, mainly Spirotrichea, were consistently detected in both fjords in spring, a pattern also observed in other Arctic fjords (Seuthe et al., 2011; Assmy et al., 2023; Bruhn et al., 2024). Ciliates are often reported to reach their highest biomass during phytoplankton spring blooms (Seuthe et al., 2011; Assmy et al., 2023), identifying them as major grazers on diatoms and bloom-associated microbes such as bacteria and flagellates (Sherr et al., 2003; Landry and Calbet, 2004). While oligotrichs and choreotrichs were also common in both fjords, loricate Tintinnina showed higher relative abundances only in inner AM during spring. They are typically described as consumers of nano-sized prey (Pierce and Turner, 1992), which aligns with the presence of such prey (mixotrophic flagellates and small green algae) in inner AM (Mikhno et al., 2026). Krawczyk et al. (Krawczyk et al., 2015, 2018) also reported generally low abundances of tintinnids in NK, except during autumn and winter, consistent with the low relative abundance of Tintinnina observed in our study. The rhizarian group Filosa-Thecofilosea was particularly abundant in NK, especially in the mid to inner fjord. The dominant group, Cryomonadida, includes taxa known to prey on diatoms (Drebes et al., 1996), so their higher importance in NK may be related to the higher diatom biomass in this fjord, especially in the inner reaches (Mikhno et al., 2026). Picozoa were important in both fjords but especially in AM, highlighting this group as a potentially pivotal component in fjord microbial food webs. To date, little is known about their role in fjord ecosystems, as most information is available for non-polar Picozoan assemblages which differ from polar ones (Huber et al., 2024). Picozoa are important grazers on picoplanktonic organisms, but may also be able to exploit sub-120 nm marine colloids as a food source (Seenivasan et al., 2013). The higher importance of Picozoa in AM may be related to the larger share of pico- and nanosized phytoplankton (Mikhno et al., 2026), but could also reflect differences in organic matter composition between the two fjord systems.

In spring, both fjords shared a broadly similar bacterial community structure dominated by Gammaproteobacteria, Bacteroidia, and Alphaproteobacteria, in line with observations from other Arctic systems (Wilson et al., 2017; De Sousa et al., 2019). Among Gammaproteobacteria, the most prominent orders were Oceanospirales—primarily uncultured *Nitrincolaceae*—alongside Cellvibrionales (notably the SAR92 clade) and Alteromonadales (*Colwellia*). These groups are generally recognized as copiotrophic responders, capable of rapidly exploiting labile organic substrates such as amino acids and polysaccharides released during phytoplankton blooms (Teeling et al., 2012, 2016; Luria et al., 2016). Although SAR92 was initially considered oligotrophic (Cho and Giovannoni, 2004), genomic evidence suggests they are able to degrade algal-derived polysaccharides such as laminarin and xylan (Xue et al., 2021), suggesting a flexible, facultatively copiotrophic lifestyle. Bacteroidia, especially Flavobacteriales (*Flavobacterium*), are effective degraders of high-molecular-weight (HMW) compounds, such as diatom-derived polysaccharides, indicating an active role in recycling phytoplankton detritus or exudates (Kirchman, 2002; Fernández-Gómez et al., 2013). Notably, *Colwellia* and *Flavobacterium* are frequently reported as dominant genera in sea-ice communities, and their relatively higher abundance in NK—particularly in the inner fjord—may be related to ice mélange and sea-ice in this area (Boetius et al., 2015). Within the Alphaproteobacteria, Rhodobacterales—particularly *Sulfitobacter*—were also abundant. They are frequently associated with phytoplankton blooms, being specialized in degrading algal-derived compounds like dimethylsulfoniopropionate (DMSP) and known to form close interactions with phytoplankton hosts (Buchan et al., 2005; Amin et al., 2015). While *Sulfitobacter* and other Rhodobacterales are traditionally recognized for their DMSP utilization, recent work in Arctic waters suggests that SAR11 (represented by *Candidatus Pelagibacter*) also correlates with DMSP catabolic genes, highlighting a broader role of this lineage in sulfur cycling than previously appreciated (Han et al., 2021). The presence of specific bacterial communities associated with the phytoplankton spring bloom is consistent with recent findings from the West Spitsbergen Current (WSC), where Priest et al. (2025) identified a recurrent spring microbial module (M3) characterized by a dominance of copiotrophic Gammaproteobacteria (including *Nitrincolaceae* and SAR92), Bacteroidia (notably Flavobacteriales), and SAR11 Clade Ia. This seasonal module was functionally enriched in genes related to carbohydrate metabolism, organic sulfur compound utilization (including DMSP catabolism), and responses to phytoplankton-derived substrates. Notably, the overlap between the M3 taxa and those observed in our dataset but also from other Arctic regions (Priest et al., 2025) supports the existence of a season-specific, bloom-associated microbiome in Arctic waters. Overall, bacterial (∼HP) abundances in spring remained relatively low in both fjords, consistent with values previously reported for the same region and season by Meire et al. (2023). Likewise, biomass of nano-sized heterotrophic protists generally remained relatively low.

### 4.2 Glacier type underlies marked restructuring and enhanced divergence in heterotrophic community composition between fjords in summer

In summer, diatom dominance in the phytoplankton decreases, except in parts of the MTG-impacted fjord NK (see below, Mikhno et al., 2026). There is a marked increase in bacterial and nano-heterotroph biomass in both fjords, especially in AM. While parasitic dinoflagellates (Syndiniales, Fig. S6) become relatively more important, rhizarian groups and Picozoa decrease. Overall, Alpha- and Gammaproteobacteria, but not Bacteroidea, remain important in both fjords, but with different specific groups, suggesting a shift towards more resource-depleted environments. In addition, differences in heterotrophic community composition and biomass between fjords became much more pronounced during summer. The seasonal restructuring and the enhanced divergence between fjords is most likely related to the increased but differential impact of glacial melt in both fjords. This leads to stronger water column stratification, surface nutrient depletion, and elevated turbidity (Mikhno et al., 2026) and a shift towards dominance of picophytoplankton (e.g. prasinophytes), heterotrophic flagellates, and dinoflagellates (Quigg et al., 2013; Dąbrowska et al., 2021; Stuart-Lee et al., 2023; Mikhno et al., 2026), especially in the LTG-impacted fjord AM (Meire et al., 2023; Stuart-Lee et al., 2023). This in turn drives a shift to increased biomass and importance of pico- and nanosized heterotrophs, most likely related to a strengthening of microbial loop interactions and recycling processes in stratified surface waters. In contrast, in NK, subsurface upwelling led to persistence of diatom growth in the mid to inner regions (Mikhno et al., 2026), with associated bacterial and heterotrophic protist (see below).

Across both fjords, *Gyrodinium* remained dominant, but other dinoflagellates, namely *Heterocapsa*, *Gymnodinium*, and *Alexandrium* increased compared to spring. *Heterocapsa* was abundant throughout AM but remained scarce in NK, whereas *Alexandrium* showed a more localized distribution. Both *Heterocapsa* and *Alexandrium* are typical peridinin-containing dinoflagellates (Zapata et al., 2012; Qiu et al., 2023). Their summer occurrence was confirmed by pigment analyses (Mikhno et al., 2026), with peridinin being detected only in AM (*Heterocapsa*) and outer NK (*Alexandrium*). The higher abundance of *Heterocapsa* in AM is consistent with the more turbid and nutrient-depleted conditions in this fjord (Mikhno et al., 2026), where its mixotrophic potential (Legrand et al., 1998; Millette et al., 2017) may provide an advantage, as also observed in other LTG Arctic fjords (Dąbrowska et al., 2021). FlowCAM analyses identified *Tripos* and to a lesser degree *Protoperidinium* as recurrent constituents of the larger microplankton assemblage. *Protoperidinium* was previously recorded from outer NK, but mainly in autumn (Krawczyk et al., 2015). These larger taxa are likely underrepresented in our metabarcoding dataset due to prefiltration steps applied prior to DNA extraction, underscoring the importance of using complementary methods (imaging-based and molecular) to fully capture community structure. Ciliates, mainly represented by Choreotrichida, persisted in outer-mid AM into summer, whereas they were comparatively scarce in both outer and inner NK. In contrast, Tintinnina significantly increased from spring to summer, representing a relatively dominant component of the larger size fraction (100–300 µm) in AM. The lower abundance of aloricate ciliates in NK during summer compared to spring may reflect increased grazing pressure by mesozooplankton, which are typically more abundant during this period (Arendt et al., 2013; Stuart-Lee et al., 2024). Notably, copepod communities in NK tend to be dominated by larger-bodied species, particularly in July (Stuart-Lee et al., 2024), which have been reported to graze on microzooplankton such as dinoflagellates and ciliates, in some cases even showing higher grazing pressure on this prey than on diatoms (Saiz and Calbet, 2011). Experimental studies further indicate that copepods often avoid tintinnids, instead selectively feeding on aloricate ciliates and heterotrophic dinoflagellates (Bouley and Kimmerer, 2006; Olson et al., 2006). Field-based observations suggest that copepod–tintinnid interactions can nevertheless vary depending on copepod species and environmental context (Rollwagen Bollens and Penry, 2003). In addition, the lower relative abundance of ciliates in the inner parts of both fjords compared to the mid stations may be linked to elevated turbidity associated with summer glacial melt. High loads of fine sediments (as evidence in the FlowCam analyses, fig. 3) can impair filter-feeding ciliates by clogging feeding structures or by adhering to loricae, increasing cell density and enhancing sinking losses. Such sediment-driven constraints on ciliates, coupled with a relative increase in dinoflagellates in more turbid fjord waters during summer, have been previously reported in Arctic fjords (Kubiszyn et al., 2014), and may represent a common response to turbid meltwater-influenced conditions.

Summer communities were characterized by a markedly higher relative abundances of Syndiniales, a widespread and diverse group of marine dinoflagellate parasites (Guillou et al., 2008; De Vargas et al., 2015) (including in Arctic waters, Lovejoy et al., 2006), which can infect a broad range of hosts— including other dinoflagellates, diatoms, ciliates, radiolarians, and metazoans—and are thus capable of top-down control on phytoplankton that is comparable to grazing (Yih and Coats, 2000; Guillou et al., 2008; Skovgaard, 2014; Sassenhagen et al., 2020). Host associations within Syndiniales vary among major clades: Group I has been linked to multiple host types, including ciliates and potentially diatoms (Skovgaard, 2014; Sassenhagen et al., 2020), whereas Group II shows a stronger and more specific association with dinoflagellates (Siano et al., 2011; Anderson and Harvey, 2020). Interpretations of host-parasite associations based on metabarcoding data alone however are challenging, as such data do not allow discrimination among distinct parasitic life stages—such as free-living spores (3–10 µm) and host-associated infective stages. By lysing host cells, Syndiniales contribute to the release of labile organic matter that supports heterotrophic bacteria and enhances microbial trophic complexity (Siano et al., 2011; Park et al., 2013). Their frequent association with sinking particles further suggests a role in vertical carbon export, linking parasitic interactions to biogeochemical fluxes (Boeuf et al., 2019; Durkin et al., 2022; Valencia et al., 2022). During summer, the highest relative abundance of Syndiniales was most pronounced below 20 m depth, consistent with conditions favouring regenerated production and intensified microbial recycling, as reflected by elevated NH₄⁺ concentrations.

In summer, while Gamma- and Alphaproteobacteria remained overall dominant, different groups became more important. The Alphaproteobacterial clade SAR11, and in particular Clade Ia, became the most abundant lineage throughout the water column (in spring it was mainly important in deeper water layers). Members of the SAR11 clade are recognized as indicators of oligotrophic marine environments (Morris et al., 2002; Giovannoni, 2017). The Gammaproteobacterial clade SAR86 increased, especially in outer-mid NK and AM. This is a globally abundant lineage with streamlined genomes adapted to low-nutrient environments, with transporters for small organic compounds and proteorhodopsin for photoheterotrophy (Dupont et al., 2012). Although typically less prominent in polar regions, SAR86 has been observed in Arctic waters during late summer, suggesting the existence of regional ecotypes adapted to cold, post-bloom conditions (Hoarfrost et al., 2020). Its prevalence in these fjords may indicate the prominence of recycled, low-molecular-weight DOM and reduced competition from copiotrophs under nutrient-poor summer stratification. Bacteroidia, particularly Flavobacteriales (e.g., *Flavobacterium* and *Polaribacter*), were consistently present but showed a marked decline in relative abundance compared to spring, especially in AM. Their persistence in NK may reflect the continued presence of diatoms in this fjord in summer. The summer increase of *Saprospiraceae* (order Chitinophagales) in AM and outer-mid NK may reflect a shift toward the degradation of more refractory organic matter, as members of this family encode diverse peptidases and carbohydrate-active enzymes (McIlroy and Nielsen, 2014; Rosenberg, 2014). Their frequent association with particle-attached niches and biofilms suggests they can exploit polymer-rich microenvironments, potentially including terrigenous particulates delivered by terrestrial runoff and meltwater (Paulsen et al., 2019). Jain et al. (2020) identified *Saprospiraceae* as one of main particle-associated taxa in Kongsfjorden, Svalbard, and demonstrated their responsiveness to complex organic substrates in fjord environments.

### 4.3 A unique ice mélange associated heterotrophic community characterizes the inner reaches of a MTG dominated fjord

In summer, the inner stations (GF10-12) of NK not only harboured unique phytoplankton groups (the haptophyte *Pseudohaptolina*, Mikhno et al., 2026) but also specific bacterial and heterotrophic protist communities that show signals of supraglacial assemblages but also of offshore, sea-ice associated communities. While in spring bacterial communities were overall similar to other parts of the fjord (and the LTG fjord), relative abundances of Gammaproteobacteria (Alteromonadales and Oceanospirales) were higher and those of Alphaproteobacteria lower. This is likely related to the more intense diatom spring bloom and enhanced availability of labile organic matter in this area (Mikhno et al., 2026), resulting in more copiotrophic conditions. Although DOM was not measured, the bacterial composition and elevated biomass suggest an active recycling of bloom-derived material. In summer, a marked shift in community composition occurred in inner NK, likely driven by intense glacial meltwater influence (e. g. iceberg melt and supraglacial runoff), which substantially modifies temperature, salinity, turbidity, and organic matter composition (Meire et al., 2015; Paulsen et al., 2019; Hopwood et al., 2020; Kellerman et al., 2020). At 1 m depth in inner NK (GF12), Gammaproteobacteria became highly dominant, with *Burkholderiaceae* alone accounting for up to 70% of the community. Within this family, *Polaromonas* was especially abundant. This genus is known for its psychrotolerance, metabolic flexibility, and adaptation to glacially influenced, low-salinity, particle-rich environments (Darcy et al., 2011; Wang et al., 2014; Gawor et al., 2016). Rassner et al. (2024) conducted a comprehensive characterization of microbial communities within the weathering crust, associated water and snow habitats on a High Arctic glacier in Svalbard, and found that *Burkholderiaceae*, in particularly *Polaromonas*, are dominant members of supraglacial assemblages. The convergence in community composition between inner NK surface water samples and glacier surface habitats suggests a direct glacial influence on the fjord assemblages. Alteromonadales—particularly *Glaciecola* and *Paraglaciecola*—also increased in this area. These genera are known copiotrophs, typically associated with the degradation of HMW-DOM during phytoplankton blooms (Teeling et al., 2012; Landa et al., 2013). Their enrichment in GF10–GF12 occurred under near-freezing conditions (∼0 °C; Mikhno et al., 2026), a niche where *Glaciecola* has been shown to thrive and to form tight associations with diatoms during cold-water blooms (Von Scheibner et al., 2017). Although local diatom production likely represents an important source of labile organic matter, glacier-derived inputs may also contribute. In particular, experiments with permafrost-derived DOM revealed that *Glaciecola* responds positively to this substrate (Müller et al., 2018). Previous studies have likewise demonstrated that glacial runoff supplies bioavailable DOM of microbial origin to downstream aquatic ecosystems (Hood et al., 2009; Bhatia et al., 2013; Christner et al., 2014; Lawson et al., 2014), potentially sustaining copiotrophic populations in addition to *in situ* sources. However, this mechanism may not be universal: in some Arctic fjords, meltwater can be relatively DOM-poor and primarily acts as a diluting agent. For example, in Young Sound (NE Greenland), visible humic-like fluorescent DOM was largely driven by the intrusion of terrestrial DOM from shelf waters and was diluted in surface waters by glacial runoff, whereas the more labile (amino acid–like) component covaried with bacterial activity and grazing, pointing to strong *in situ* transformation pathways (Paulsen et al., 2019). These contrasting patterns highlight the need for a more comprehensive characterization of DOM sources and transformations in Arctic fjords, which are influenced not only by glacier type, but also by *in situ* productivity, coastal water inputs, and fjord circulation dynamics. Verrucomicrobiae, in particular *Lentimonas*, increased in abundance during summer, particularly in the inner NK, suggesting an adaptation to environments enriched in complex polysaccharides (Spring et al., 2016). Similar patterns have been observed in Svalbard fjords, where members of this genus were implicated in the degradation of specific HMW polysaccharides (Cardman et al., 2014). Finally, in inner NK surface waters, the SAR11 clade nearly vanished (< 1%). The observed positive correlation between SAR11 Clade Ia and temperature suggests that the markedly colder conditions in the inner fjord may constrain growth of this lineage. This pattern, together with the shift toward a clearly more copiotrophic bacterial community, indicates environmental conditions under which oligotrophic specialists such as SAR11 may become less competitive.

### 4.4 Deep-water communities

At depth (≥ 150 m), microbial communities showed limited seasonal restructuring compared with surface waters (1–40 m), but deeper layers became overall more distinct in summer. Dinophyceae dominated deep-water communities in spring, with their relative abundance decreasing up-fjord, where Syndiniales—particularly Dino-Group II—became more prominent, suggesting an increase in parasitic interactions that may have disadvantaged dominant Dinophyceae. In summer, Syndiniales became overall dominant at depth, emphasizing the role of parasitism and particle-associated recycling in sustaining deep-water microbial activity (Boeuf et al., 2019; Preston et al., 2020; Cruz et al., 2021; Anderson et al., 2024). Anderson et al. (2024) showed a significant negative relationship between the abundance of Syndiniales in surface waters and the flux of particulate organic carbon (POC) at depth, suggesting that parasitic activity may reduce carbon export by rerouting host-derived carbon toward dissolved and small particulate pools, thereby enhancing remineralization and attenuating particle flux. In the context of fjords, which act as sites of organic carbon burial (Smith et al., 2015), these processes may have important implications for carbon export and warrant further investigation. In both seasons, bacterial communities at depth were dominated by SAR11 Clade Ia. This pattern is noteworthy, as this clade is typically considered a surface-associated ecotype and generally declines below the euphotic zone (Field et al., 1997; Carlson et al., 2009; Giovannoni, 2017). Its presence at depth may reflect enhanced vertical export of surface-derived material, consistent with occasional reports of this clade in mesopelagic waters under strong export conditions (Carlson et al., 2009; Thiele et al., 2023). Importantly, iFCM counts indicated that total bacterial abundances at depth were relatively low (∼5 x 10^4^ cells mL⁻¹), so the high relative abundance of Clade Ia reflects relative dominance within a low abundance bacterial community. 16S-based assignments cannot resolve the fine-scale diversity within Clade Ia, and further phylogenetic or functional analyses would be required to clarify potential ecotypic differences. Other taxa enriched in deep waters included the NS9 marine group, SAR11 Clade II, and SUP05. The occurrence of the NS9 marine group is consistent with a role in the processing of particle-derived or more complex organic matter, as members of this Flavobacteriales lineage are often associated with particulate organic matter and depth-related community differentiation in marine systems (Yeh and Fuhrman, 2022; Reintjes et al., 2023). Within SAR11, the enrichment of Clade II at depth relative to surface waters reflects well-established vertical niche partitioning among SAR11 ecotypes, with this clade typically more abundant in mesopelagic waters with low-light and low-energy conditions (Giovannoni, 2017; Bolaños et al., 2022; Hays and Fuchsman, 2025). Finally, the presence of SUP05 is noteworthy, as this clade is well known to comprise sulfur-oxidizing, chemolithotrophic Gammaproteobacteria from deep marine environments (Anantharaman et al., 2013; Marshall and Morris, 2013). Its occurrence thus suggests that part of the deep bacterial assemblage may be supported by sulfur-based energy metabolism.

### 4.5 Increased environmental filtering and ecological coupling in pelagic microbial fjord communities in summer

The variance partitioning and Mantel analyses indicate that microbial community structure in the fjords is shaped by both environmental filtering and biological interactions. Although environmental variables explained the largest independent fraction of variance, this contribution was relatively modest, with a substantial portion being shared between environmental, phytoplankton, heterotrophic protist, and bacterial components. This suggests that environmental gradients act as overarching constraints, with community variability arising from coordinated responses within interconnected microbial networks, consistent with network-based views of plankton ecosystems (Lima-Mendez et al., 2015). The marked increase in Mantel intercorrelations amongst the different microbial components and with the environmental variables in summer, but also in the shared fractions of explained community variance, suggest that environmental filtering and/or biotic interactions strongly increased in this season, especially in the MTG-dominated NK fjord. Most likely this is related to enhanced glacial melt leading to stronger and more stable water column stratification (and hence vertical niche differentiation). However, the stronger microbial intercorrelation structure may also hint at more closely coupled interactions, such as competition and facilitation, in an overall more resource depleted environment in summer.

## 5 Conclusion

Overall, this study shows that heterotrophic microbial communities in NK and AM undergo strong seasonal restructuring, with summer marked by increased pico- and nano-heterotrophic biomass, higher importance of parasitic dinoflagellates and oligotrophic bacterial lineages, and stronger differentiation between surface and deep communities. This restructuring amplified the ecological contrast between the two fjords: AM shifted toward a more recycling-dominated microbial food web under stratified, nutrient-depleted conditions, whereas NK retained stronger links to diatom production and, in its inner reaches, hosted a distinct ice- and glacier-influenced microbial assemblage. Together, these patterns indicate that glacier type and seasonal meltwater dynamics strongly shape microbial food-web organization in Arctic fjords. Future work should combine seasonal time series with measurements of DOM composition, microbial activity, and host–parasite interactions to better resolve the functional consequences of these community shifts for carbon cycling and export.

## Financial support

This work is part of the IMAGIN project, funded by Research Foundation – Flanders (FWO) (grant no. 3G043120) and by the Belgian Federal Science Policy Office (BELSPO)-funded project CANOE (contract no. RV/21/CANOE).

The research leading to the results presented in this publication was carried out with infrastructure funded by EMBRC Belgium–FWO international research infrastructures I001621N and I000825N.

## Acknowledgements

We thank the captain of the RV *Avataq* for his invaluable assistance during fieldwork. We are also grateful to the staff of the Greenland Institute of Natural Resources (GINR) and to all collaborators in Greenland for their logistical and technical support during the sampling campaigns.

## Data Availability

The dataset supporting this study has been deposited in the Zenodo repository (https://doi.org/10.5281/zenodo.21381561). The high throughput sequence data have been deposited in the NCBI SRA under BioProject accession PRJNA1457328 (eukaryotes) and PRJNA1466576 (bacteria).

## Conflict of Interest

The authors declare that the research was conducted in the absence of any commercial or financial relationships that could be construed as a potential conflict of interest.

## Notes

### Competing Interest Statement

The authors have declared no competing interest.

## References

Amin, S. A., Hmelo, L. R., Van Tol, H. M., Durham, B. P., Carlson, L. T., Heal, K. R., et al. (2015). Interaction and signalling between a cosmopolitan phytoplankton and associated bacteria. Nature 522, 98–101. doi: 10.1038/nature14488

Anantharaman, K., Breier, J. A., Sheik, C. S., and Dick, G. J. (2013). Evidence for hydrogen oxidation and metabolic plasticity in widespread deep-sea sulfur-oxidizing bacteria. Proc. Natl. Acad. Sci. U.S.A. 110, 330–335. doi: 10.1073/pnas.1215340110

Anderson, M. J., and Willis, T. J. (2003). CANONICAL ANALYSIS OF PRINCIPAL COORDINATES: A USEFUL METHOD OF CONSTRAINED ORDINATION FOR ECOLOGY. Ecology 84, 511–525. doi: 10.1890/0012-9658(2003)084[0511:CAOPCA]2.0.CO;2

Anderson, S. R., Blanco-Bercial, L., Carlson, C. A., and Harvey, E. L. (2024). Role of Syndiniales parasites in depth-specific networks and carbon flux in the oligotrophic ocean. ISME Communications 4, ycae014. doi: 10.1093/ismeco/ycae014

Anderson, S. R., and Harvey, E. L. (2020). Temporal Variability and Ecological Interactions of Parasitic Marine Syndiniales in Coastal Protist Communities. mSphere 5, e00209–20. doi: 10.1128/mSphere.00209-20

Andresen, C. S., Karlsson, N. B., Straneo, F., Schmidt, S., Andersen, T. J., Eidam, E. F., et al. (2024). Sediment discharge from Greenland’s marine-terminating glaciers is linked with surface melt. Nat Commun 15, 1332. doi: 10.1038/s41467-024-45694-1

Arendt, K. E., Juul-Pedersen, T., Mortensen, J., Blicher, M. E., and Rysgaard, S. (2013). A 5-year study of seasonal patterns in mesozooplankton community structure in a sub-Arctic fjord reveals dominance of Microsetella norvegica (Crustacea, Copepoda). Journal of Plankton Research 35, 105–120. doi: 10.1093/plankt/fbs087

Assmy, P., Cecilie Kvernvik, A., Hop, H., Hoppe, C. J. M., Chierici, M., David T., D., et al. (2023). Seasonal plankton dynamics in Kongsfjorden during two years of contrasting environmental conditions. Progress in Oceanography 213, 102996. doi: 10.1016/j.pocean.2023.102996

Azam, F., Fenchel, T., Field, J., Gray, J., Meyer-Reil, L., and Thingstad, F. (1983). The Ecological Role of Water-Column Microbes in the Sea. Mar. Ecol. Prog. Ser. 10, 257–263. doi: 10.3354/meps010257

Bhatia, M. P., Das, S. B., Xu, L., Charette, M. A., Wadham, J. L., and Kujawinski, E. B. (2013). Organic carbon export from the Greenland ice sheet. Geochimica et Cosmochimica Acta 109, 329–344. doi: 10.1016/j.gca.2013.02.006

Boetius, A., Anesio, A. M., Deming, J. W., Mikucki, J. A., and Rapp, J. Z. (2015). Microbial ecology of the cryosphere: sea ice and glacial habitats. Nat Rev Microbiol 13, 677–690. doi: 10.1038/nrmicro3522

Boeuf, D., Edwards, B. R., Eppley, J. M., Hu, S. K., Poff, K. E., Romano, A. E., et al. (2019). Biological composition and microbial dynamics of sinking particulate organic matter at abyssal depths in the oligotrophic open ocean. Proc. Natl. Acad. Sci. U.S.A. 116, 11824–11832. doi: 10.1073/pnas.1903080116

Bolaños, L. M., Tait, K., Somerfield, P. J., Parsons, R. J., Giovannoni, S. J., Smyth, T., et al. (2022). Influence of short and long term processes on SAR11 communities in open ocean and coastal systems. ISME Communications 2, 116. doi: 10.1038/s43705-022-00198-1

Bouley, P., and Kimmerer, W. (2006). Ecology of a highly abundant, introduced cyclopoid copepod in a temperate estuary. Mar. Ecol. Prog. Ser. 324, 219–228. doi: 10.3354/meps324219

Bruhn, C. S., Lundholm, N., Hansen, P. J., Wohlrab, S., and John, U. (2024). Transition from a mixotrophic/heterotrophic protist community during the dark winter to a photoautotrophic spring community in surface waters of Disko Bay, Greenland. Front. Microbiol. 15, 1407888. doi: 10.3389/fmicb.2024.1407888

Buchan, A., González, J. M., and Moran, M. A. (2005). Overview of the Marine *Roseobacter* Lineage. Appl Environ Microbiol 71, 5665–5677. doi: 10.1128/AEM.71.10.5665-5677.2005

Calbet, A., and Landry, M. R. (2004). Phytoplankton growth, microzooplankton grazing, and carbon cycling in marine systems. Limnology & Oceanography 49, 51–57. doi: 10.4319/lo.2004.49.1.0051

Callahan, B. J., McMurdie, P. J., Rosen, M. J., Han, A. W., Johnson, A. J. A., and Holmes, S. P. (2016). DADA2: High-resolution sample inference from Illumina amplicon data. Nat Methods 13, 581–583. doi: 10.1038/nmeth.3869

Cameron, K. A., Stibal, M., Hawkings, J. R., Mikkelsen, A. B., Telling, J., Kohler, T. J., et al. (2017). Meltwater export of prokaryotic cells from the Greenland ice sheet. Environmental Microbiology 19, 524–534. doi: 10.1111/1462-2920.13483

Cape, M. R., Straneo, F., Beaird, N., Bundy, R. M., and Charette, M. A. (2019). Nutrient release to oceans from buoyancy-driven upwelling at Greenland tidewater glaciers. Nature Geosci 12, 34–39. doi: 10.1038/s41561-018-0268-4

Cardman, Z., Arnosti, C., Durbin, A., Ziervogel, K., Cox, C., Steen, A. D., et al. (2014). Verrucomicrobia Are Candidates for Polysaccharide-Degrading Bacterioplankton in an Arctic Fjord of Svalbard. Appl Environ Microbiol 80, 3749–3756. doi: 10.1128/AEM.00899-14

Carlson, C. A., Morris, R., Parsons, R., Treusch, A. H., Giovannoni, S. J., and Vergin, K. (2009). Seasonal dynamics of SAR11 populations in the euphotic and mesopelagic zones of the northwestern Sargasso Sea. The ISME Journal 3, 283–295. doi: 10.1038/ismej.2008.117

Catania, G. A., Stearns, L. A., Moon, T. A., Enderlin, E. M., and Jackson, R. H. (2020). Future Evolution of Greenland’s Marine-Terminating Outlet Glaciers. JGR Earth Surface 125, e2018JF004873. doi: 10.1029/2018JF004873

Cho, J.-C., and Giovannoni, S. J. (2004). Cultivation and Growth Characteristics of a Diverse Group of Oligotrophic Marine *Gammaproteobacteria*. Appl Environ Microbiol 70, 432–440. doi: 10.1128/AEM.70.1.432-440.2004

Christner, B. C., Priscu, J. C., Achberger, A. M., Barbante, C., Carter, S. P., Christianson, K., et al. (2014). A microbial ecosystem beneath the West Antarctic ice sheet. Nature 512, 310–313. doi: 10.1038/nature13667

Chua, S. D. X., Mortensen, J., Uotila, P., and Meire, L. (2025). Influence of Ocean Waters in Retreat Episodes of a West Greenland Tidewater Outlet Glacier. JGR Oceans 130, e2024JC022161. doi: 10.1029/2024JC022161

Cruz, B. N., Brozak, S., and Neuer, S. (2021). Microscopy and DNA -based characterization of sinking particles at the Bermuda Atlantic Time-series Study station point to zooplankton mediation of particle flux. Limnology & Oceanography 66, 3697–3713. doi: 10.1002/lno.11910

Dąbrowska, A. M., Wiktor, J. M., Wiktor, J. M., Kristiansen, S., Vader, A., and Gabrielsen, T. (2021). When a Year Is Not Enough: Further Study of the Seasonality of Planktonic Protist Communities Structure in an Ice-Free High Arctic Fjord (Adventfjorden, West Spitsbergen). Water 13, 1990. doi: 10.3390/w13141990

Darcy, J. L., Lynch, R. C., King, A. J., Robeson, M. S., and Schmidt, S. K. (2011). Global Distribution of Polaromonas Phylotypes - Evidence for a Highly Successful Dispersal Capacity. PLoS ONE 6, e23742. doi: 10.1371/journal.pone.0023742

De Coster, W., D’Hert, S., Schultz, D. T., Cruts, M., and Van Broeckhoven, C. (2018). NanoPack: visualizing and processing long-read sequencing data. Bioinformatics 34, 2666–2669. doi: 10.1093/bioinformatics/bty149

De Sousa, A. G. G., Tomasino, M. P., Duarte, P., Fernández-Méndez, M., Assmy, P., Ribeiro, H., et al. (2019). Diversity and Composition of Pelagic Prokaryotic and Protist Communities in a Thin Arctic Sea-Ice Regime. Microb Ecol 78, 388–408. doi: 10.1007/s00248-018-01314-2

De Vargas, C., Audic, S., Henry, N., Decelle, J., Mahé, F., Logares, R., et al. (2015). Eukaryotic plankton diversity in the sunlit ocean. Science 348, 1261605. doi: 10.1126/science.1261605

Drebes, G., Kühn, S. F., Gmelch, A., and Schnepf, E. (1996). Cryothecomonas aestivalis sp. nov., a colourless nanoflagellate feeding on the marine centric diatomGuinardia delicatula (Cleve) Hasle. Helgolander Meeresunters 50, 497–515. doi: 10.1007/BF02367163

Dupont, C. L., Rusch, D. B., Yooseph, S., Lombardo, M.-J., Alexander Richter, R., Valas, R., et al. (2012). Genomic insights to SAR86, an abundant and uncultivated marine bacterial lineage. The ISME Journal 6, 1186–1199. doi: 10.1038/ismej.2011.189

Durkin, C. A., Cetinić, I., Estapa, M., Ljubešić, Z., Mucko, M., Neeley, A., et al. (2022). Tracing the path of carbon export in the ocean though DNA sequencing of individual sinking particles. The ISME Journal 16, 1896–1906. doi: 10.1038/s41396-022-01239-2

Engel Arendt, K., Nielsen, T., Rysgaard, S., and Tönnesson, K. (2010). Differences in plankton community structure along the Godthåbsfjord, from the Greenland Ice Sheet to offshore waters. Mar. Ecol. Prog. Ser. 401, 49–62. doi: 10.3354/meps08368

European Space Agency (ESA) (2025). Copernicus Sentinel-2 data. Sentinel Hub EO Browser. Available at: https://apps.sentinel-hub.com/eo-browser/

Fernández-Gómez, B., Richter, M., Schüler, M., Pinhassi, J., Acinas, S. G., González, J. M., et al. (2013). Ecology of marine Bacteroidetes: a comparative genomics approach. The ISME Journal 7, 1026–1037. doi: 10.1038/ismej.2012.169

Field, K. G., Gordon, D., Wright, T., Rappé, M., Urback, E., Vergin, K., et al. (1997). Diversity and depth-specific distribution of SAR11 cluster rRNA genes from marine planktonic bacteria. Appl Environ Microbiol 63, 63–70. doi: 10.1128/aem.63.1.63-70.1997

Gawor, J., Grzesiak, J., Sasin-Kurowska, J., Borsuk, P., Gromadka, R., Górniak, D., et al. (2016). Evidence of adaptation, niche separation and microevolution within the genus Polaromonas on Arctic and Antarctic glacial surfaces. Extremophiles 20, 403–413. doi: 10.1007/s00792-016-0831-0

Giovannoni, S. J. (2017). SAR11 Bacteria: The Most Abundant Plankton in the Oceans. Annu. Rev. Mar. Sci. 9, 231–255. doi: 10.1146/annurev-marine-010814-015934

Graßhoff, K., Graßhoff, K., Kremling, K., and Ehrhardt, M. eds. (2009). Methods of Seawater Analysis., 3. vollst. überarb. u. erw. Auflage. Weinheim: Wiley-VCH.

Guillou, L., Viprey, M., Chambouvet, A., Welsh, R. M., Kirkham, A. R., Massana, R., et al. (2008). Widespread occurrence and genetic diversity of marine parasitoids belonging to *Syndiniales* ( *Alveolata*). Environmental Microbiology 10, 3349–3365. doi: 10.1111/j.1462-2920.2008.01731.x

Han, D., Park, K.-T., Kim, H., Kim, T.-H., Jeong, M.-K., and Nam, S.-I. (2024). Interaction between phytoplankton and heterotrophic bacteria in Arctic fjords during the glacial melting season as revealed by eDNA metabarcoding. FEMS Microbiology Ecology 100, fiae059. doi: 10.1093/femsec/fiae059

Han, D., Richter-Heitmann, T., Kim, I.-N., Choy, E., Park, K.-T., Unno, T., et al. (2021). Survey of Bacterial Phylogenetic Diversity During the Glacier Melting Season in an Arctic Fjord. Microb Ecol 81, 579–591. doi: 10.1007/s00248-020-01616-4

Hays, M. D., and Fuchsman, C. A. (2025). SAR11 ecotypes across ocean basins change with depth due to changes in light and oxygen. The ISME Journal 19, wraf221. doi: 10.1093/ismejo/wraf221

Hoarfrost, A., Nayfach, S., Ladau, J., Yooseph, S., Arnosti, C., Dupont, C. L., et al. (2020). Global ecotypes in the ubiquitous marine clade SAR86. The ISME Journal 14, 178–188. doi: 10.1038/s41396-019-0516-7

Hood, E., Fellman, J., Spencer, R. G. M., Hernes, P. J., Edwards, R., D’Amore, D., et al. (2009). Glaciers as a source of ancient and labile organic matter to the marine environment. Nature 462, 1044– 1047. doi: 10.1038/nature08580

Hopwood, M. J., Carroll, D., Browning, T. J., Meire, L., Mortensen, J., Krisch, S., et al. (2018). Non-linear response of summertime marine productivity to increased meltwater discharge around Greenland. Nat Commun 9, 3256. doi: 10.1038/s41467-018-05488-8

Hopwood, M. J., Carroll, D., Dunse, T., Hodson, A., Holding, J. M., Iriarte, J. L., et al. (2020). Review article: How does glacier discharge affect marine biogeochemistry and primary production in the Arctic? The Cryosphere 14, 1347–1383. doi: 10.5194/tc-14-1347-2020

Huber, P., De Angelis, D., Sarmento, H., Metz, S., Giner, C. R., Vargas, C. D., et al. (2024). Global distribution, diversity, and ecological niche of Picozoa, a widespread and enigmatic marine protist lineage. Microbiome 12, 162. doi: 10.1186/s40168-024-01874-1

Jain, A., Krishnan, K. P., Begum, N., Singh, A., Thomas, F. A., and Gopinath, A. (2020). Response of bacterial communities from Kongsfjorden (Svalbard, Arctic Ocean) to macroalgal polysaccharide amendments. Marine Environmental Research 155, 104874. doi: 10.1016/j.marenvres.2020.104874

Joughin, I., Smith, B. E., Howat, I. M., Scambos, T., and Moon, T. (2010). Greenland flow variability from ice-sheet-wide velocity mapping. J. Glaciol. 56, 415–430. doi: 10.3189/002214310792447734

Juul-Pedersen, T., Arendt, K., Mortensen, J., Blicher, M., Søgaard, D., and Rysgaard, S. (2015). Seasonal and interannual phytoplankton production in a sub-Arctic tidewater outlet glacier fjord, SW Greenland. Mar. Ecol. Prog. Ser. 524, 27–38. doi: 10.3354/meps11174

Kavan, J., Szczypińska, M., Kochtitzky, W., Farquharson, L., Bendixen, M., and Strzelecki, M. C. (2025). New coasts emerging from the retreat of Northern Hemisphere marine-terminating glaciers in the twenty-first century. Nat. Clim. Chang. 15, 528–537. doi: 10.1038/s41558-025-02282-5

Kellerman, A. M., Hawkings, J. R., Wadham, J. L., Kohler, T. J., Stibal, M., Grater, E., et al. (2020). Glacier Outflow Dissolved Organic Matter as a Window Into Seasonally Changing Carbon Sources: Leverett Glacier, Greenland. JGR Biogeosciences 125, e2019JG005161. doi: 10.1029/2019JG005161

Kirchman, D. (2002). The ecology of Cytophaga–Flavobacteria in aquatic environments. FEMS Microbiology Ecology 39, 91–100. doi: 10.1016/S0168-6496(01)00206-9

Krawczyk, D. W., Meire, L., Lopes, C., Juul-Pedersen, T., Mortensen, J., Li, C. L., et al. (2018). Seasonal succession, distribution, and diversity of planktonic protists in relation to hydrography of the Godthåbsfjord system (SW Greenland). Polar Biol 41, 2033–2052. doi: 10.1007/s00300-018-2343-0

Krawczyk, D. W., Witkowski, A., Juul-Pedersen, T., Arendt, K. E., Mortensen, J., and Rysgaard, S. (2015). Microplankton succession in a SW Greenland tidewater glacial fjord influenced by coastal inflows and run-off from the Greenland Ice Sheet. Polar Biol 38, 1515–1533. doi: 10.1007/s00300-015-1715-y

Kubiszyn, A. M., Piwosz, K., Wiktor, J. M., and Wiktor, J. M. (2014). The effect of inter-annual Atlantic water inflow variability on the planktonic protist community structure in the West Spitsbergen waters during the summer. Journal of Plankton Research 36, 1190–1203. doi: 10.1093/plankt/fbu044

Landa, M., Cottrell, M., Kirchman, D., Blain, S., and Obernosterer, I. (2013). Changes in bacterial diversity in response to dissolved organic matter supply in a continuous culture experiment. Aquat. Microb. Ecol. 69, 157–168. doi: 10.3354/ame01632

Landry, M. R., and Calbet, A. (2004). Microzooplankton production in the oceans. ICES Journal of Marine Science 61, 501–507. doi: 10.1016/j.icesjms.2004.03.011

Lawson, E. C., Wadham, J. L., Tranter, M., Stibal, M., Lis, G. P., Butler, C. E. H., et al. (2014). Greenland Ice Sheet exports labile organic carbon to the Arctic oceans. Biogeosciences 11, 4015–4028. doi: 10.5194/bg-11-4015-2014

Legrand, C., Granéli, E., and Carlsson, P. (1998). Induced phagotrophy in the photosynthetic dinoflagellate Heterocapsa triquetra. Aquat. Microb. Ecol. 15, 65–75. doi: 10.3354/ame015065

Lima-Mendez, G., Faust, K., Henry, N., Decelle, J., Colin, S., Carcillo, F., et al. (2015). Determinants of community structure in the global plankton interactome. Science 348, 1262073. doi: 10.1126/science.1262073

Lin, S. (2006). THE SMALLEST DINOFLAGELLATE GENOME IS YET TO BE FOUND: A COMMENT ON LAJEUNESSE ET AL. “ *SYMBIODINIUM* (PYRRHOPHYTA) GENOME SIZES (DNA CONTENT) ARE SMALLEST AMONG DINOFLAGELLATES”^1^. Journal of Phycology 42, 746–748. doi: 10.1111/j.1529-8817.2006.00213.x

Lovejoy, C., Massana, R., and Pedrós-Alió, C. (2006). Diversity and Distribution of Marine Microbial Eukaryotes in the Arctic Ocean and Adjacent Seas. Appl Environ Microbiol 72, 3085–3095. doi: 10.1128/AEM.72.5.3085-3095.2006

Lu, J., and Salzberg, S. L. (2020). Ultrafast and accurate 16S rRNA microbial community analysis using Kraken 2. Microbiome 8, 124. doi: 10.1186/s40168-020-00900-2

Paulsen, M. L., Müller, O., Larsen, A., Møller, E. F., Middelboe, M., Sejr, M. K., et al. (2019). Biological transformation of Arctic dissolved organic matter in a NE Greenland fjord. Limnology & Oceanography 64, 1014–1033. doi: 10.1002/lno.11091

Luria, C. M., Amaral-Zettler, L. A., Ducklow, H. W., and Rich, J. J. (2016). Seasonal Succession of Free-Living Bacterial Communities in Coastal Waters of the Western Antarctic Peninsula. Front. Microbiol. 7. doi: 10.3389/fmicb.2016.01731

Marijon, P., Chikhi, R., and Varré, J.-S. (2020). yacrd and fpa: upstream tools for long-read genome assembly. Bioinformatics 36, 3894–3896. doi: 10.1093/bioinformatics/btaa262

Marshall, K. T., and Morris, R. M. (2013). Isolation of an aerobic sulfur oxidizer from the SUP05/Arctic96BD-19 clade. The ISME Journal 7, 452–455. doi: 10.1038/ismej.2012.78

McIlroy, S. J., and Nielsen, P. H. (2014). “The Family Saprospiraceae,” in The Prokaryotes, eds. E. Rosenberg, E. F. DeLong, S. Lory, E. Stackebrandt, and F. Thompson (Berlin, Heidelberg: Springer Berlin Heidelberg), 863–889. doi: 10.1007/978-3-642-38954-2_138

McMurdie, P. J., and Holmes, S. (2013). phyloseq: An R Package for Reproducible Interactive Analysis and Graphics of Microbiome Census Data. PLoS ONE 8, e61217. doi: 10.1371/journal.pone.0061217

Meire, L., Meire, P., Struyf, E., Krawczyk, D. W., Arendt, K. E., Yde, J. C., et al. (2016a). High export of dissolved silica from the Greenland Ice Sheet. Geophysical Research Letters 43, 9173–9182. doi: 10.1002/2016GL070191

Meire, L., Mortensen, J., Meire, P., Juul-Pedersen, T., Sejr, M. K., Rysgaard, S., et al. (2017). Marine-terminating glaciers sustain high productivity in Greenland fjords. Global Change Biology 23, 5344–5357. doi: 10.1111/gcb.13801

Meire, L., Mortensen, J., Rysgaard, S., Bendtsen, J., Boone, W., Meire, P., et al. (2016b). Spring bloom dynamics in a subarctic fjord influenced by tidewater outlet glaciers (Godthåbsfjord, SW Greenland). JGR Biogeosciences 121, 1581–1592. doi: 10.1002/2015JG003240

Meire, L., Paulsen, M. L., Meire, P., Rysgaard, S., Hopwood, M. J., Sejr, M. K., et al. (2023). Glacier retreat alters downstream fjord ecosystem structure and function in Greenland. Nat. Geosci. 16, 671–674. doi: 10.1038/s41561-023-01218-y

Meire, L., Søgaard, D. H., Mortensen, J., Meysman, F. J. R., Soetaert, K., Arendt, K. E., et al. (2015). Glacial meltwater and primary production are drivers of strong CO_2_ uptake in fjord and coastal waters adjacent to the Greenland Ice Sheet. Biogeosciences 12, 2347–2363. doi: 10.5194/bg-12-2347-2015

Menden-Deuer, S., and Lessard, E. J. (2000). Carbon to volume relationships for dinoflagellates, diatoms, and other protist plankton. Limnology & Oceanography 45, 569–579. doi: 10.4319/lo.2000.45.3.0569

Middelboe, M., Glud, R. N., and Sejr, M. K. (2012). Bacterial carbon cycling in a subarctic fjord: A seasonal study on microbial activity, growth efficiency, and virus-induced mortality in Kobbefjord, Greenland. Limnology & Oceanography 57, 1732–1742. doi: 10.4319/lo.2012.57.6.1732

Mikhno, M., Meire, L., Dasseville, R., Chaerle, P., Daveloose, I., D’hondt, S., et al. (2026). Seasonal divergence in phytoplankton community and size structure in contrasting fjord types in SW Greenland. [Preprint]. doi: 10.5194/egusphere-2026-5308

Millette, N. C., Pierson, J. J., Aceves, A., and Stoecker, D. K. (2017). Mixotrophy in *Heterocapsa rotundata* : A mechanism for dominating the winter phytoplankton. Limnology & Oceanography 62, 836–845. doi: 10.1002/lno.10470

Morris, R. M., Rappé, M. S., Connon, S. A., Vergin, K. L., Siebold, W. A., Carlson, C. A., et al. (2002). SAR11 clade dominates ocean surface bacterioplankton communities. Nature 420, 806–810. doi: 10.1038/nature01240

Mortensen, J., Bendtsen, J., Motyka, R. J., Lennert, K., Truffer, M., Fahnestock, M., et al. (2013). On the seasonal freshwater stratification in the proximity of fast-flowing tidewater outlet glaciers in a sub-Arctic sill fjord. JGR Oceans 118, 1382–1395. doi: 10.1002/jgrc.20134

Mortensen, J., Lennert, K., Bendtsen, J., and Rysgaard, S. (2011). Heat sources for glacial melt in a sub-Arctic fjord (Godthåbsfjord) in contact with the Greenland Ice Sheet. J. Geophys. Res. 116, C01013. doi: 10.1029/2010JC006528

Müller, O., Seuthe, L., Bratbak, G., and Paulsen, M. L. (2018). Bacterial Response to Permafrost Derived Organic Matter Input in an Arctic Fjord. Front. Mar. Sci. 5, 263. doi: 10.3389/fmars.2018.00263

Murray, T., Scharrer, K., Selmes, N., Booth, A. D., James, T. D., Bevan, S. L., et al. (2015). Extensive retreat of Greenland tidewater glaciers, 2000–2010. *Arctic, Antarctic, and Alpine Research* 47, 427–447. doi: 10.1657/AAAR0014-049

Oksanen, J., Simpson, G. L., Blanchet, F. G., Kindt, R., Legendre, P., Minchin, P. R., et al. (2025). vegan: Community Ecology Package. 2.7-5. doi: 10.32614/CRAN.package.vegan

Oksman, M., Kvorning, A. B., Larsen, S. H., Kjeldsen, K. K., Mankoff, K. D., Colgan, W., et al. (2022). Impact of freshwater runoff from the southwest Greenland Ice Sheet on fjord productivity since the late 19th century. The Cryosphere 16, 2471–2491. doi: 10.5194/tc-16-2471-2022

Olson, M., Lessard, E., Wong, C., and Bernhardt, M. (2006). Copepod feeding selectivity on microplankton, including the toxigenic diatoms Pseudo-nitzschia spp., in the coastal Pacific Northwest. Mar. Ecol. Prog. Ser. 326, 207–220. doi: 10.3354/meps326207

Park, M. G., Kim, S., Shin, E.-Y., Yih, W., and Coats, D. W. (2013). Parasitism of harmful dinoflagellates in Korean coastal waters. Harmful Algae 30, S62–S74. doi: 10.1016/j.hal.2013.10.007

Pierce, R. W., and Turner, J. T. (1992). Ecology of planktonic ciliates in marine food webs. 6, 139–181.

Pomeroy, L. R. (1974). The Ocean’s Food Web, A Changing Paradigm. BioScience 24, 499–504. doi: 10.2307/1296885

Preston, C. M., Durkin, C. A., and Yamahara, K. M. (2020). DNA metabarcoding reveals organisms contributing to particulate matter flux to abyssal depths in the North East Pacific ocean. Deep Sea Research Part II: Topical Studies in Oceanography 173, 104708. doi: 10.1016/j.dsr2.2019.104708

Previdi, M., Smith, K. L., and Polvani, L. M. (2021). Arctic amplification of climate change: a review of underlying mechanisms. Environ. Res. Lett. 16, 093003. doi: 10.1088/1748-9326/ac1c29

Priest, T., Oldenburg, E., Popa, O., Dede, B., Metfies, K., Von Appen, W.-J., et al. (2025). Seasonal recurrence and modular assembly of an Arctic pelagic marine microbiome. Nat Commun 16, 1326. doi: 10.1038/s41467-025-56203-3

Qiu, S., Yuan, Y., Li, X., Zhao, C., He, Y., Tang, B., et al. (2023). Peridinin-chlorophyll-protein complex industry from algae: A critical review of the current advancements, hurdles, and biotechnological potential. Algal Research 72, 103118. doi: 10.1016/j.algal.2023.103118

Quigg, A., Nunnally, C., McInnes, A., Gay, S., Rowe, G., Dellapenna, T., et al. (2013). Hydrographic and biological controls in two subarctic fjords: an environmental case study of how climate change could impact phytoplankton communities. Mar. Ecol. Prog. Ser. 480, 21–37. doi: 10.3354/meps10225

R Core Team (2021). R: A Language and Environment for Statistical Computing.

Rantanen, M., Karpechko, A. Yu., Lipponen, A., Nordling, K., Hyvärinen, O., Ruosteenoja, K., et al. (2022). The Arctic has warmed nearly four times faster than the globe since 1979. Commun Earth Environ 3, 168. doi: 10.1038/s43247-022-00498-3

Rassner, S. M. E., Cook, J. M., Mitchell, A. C., Stevens, I. T., Irvine-Fynn, T. D. L., Hodson, A. J., et al. (2024). The distinctive weathering crust habitat of a High Arctic glacier comprises discrete microbial micro-habitats. Environmental Microbiology 26, e16617. doi: 10.1111/1462-2920.16617

Reintjes, G., Heins, A., Wang, C., and Amann, R. (2023). Abundance and composition of particles and their attached microbiomes along an Atlantic Meridional Transect. Front. Mar. Sci. 10, 1051510. doi: 10.3389/fmars.2023.1051510

Rodríguez-Marconi, S., Krock, B., Tillmann, U., Tillmann, A., Voss, D., Zielinski, O., et al. (2024). Diversity of eukaryote plankton and phycotoxins along the West Kalaallit Nunaat (Greenland) coast. Front. Mar. Sci. 11, 1443389. doi: 10.3389/fmars.2024.1443389

Rokkan Iversen, K., and Seuthe, L. (2011). Seasonal microbial processes in a high-latitude fjord (Kongsfjorden, Svalbard): I. Heterotrophic bacteria, picoplankton and nanoflagellates. Polar Biol 34, 731–749. doi: 10.1007/s00300-010-0929-2

Rollwagen Bollens, G., and Penry, D. (2003). Feeding dynamics of Acartia spp. copepods in a large, temperate estuary (San Francisco Bay, CA). Mar. Ecol. Prog. Ser. 257, 139–158. doi: 10.3354/meps257139

Romanova, N. D., and Sazhin, A. F. (2010). Relationships between the cell volume and the carbon content of bacteria. Oceanology 50, 522–530. doi: 10.1134/S0001437010040089

Rosenberg, E. (2014). “The Family Chitinophagaceae,” in The Prokaryotes, eds. E. Rosenberg, E. F. DeLong, S. Lory, E. Stackebrandt, and F. Thompson (Berlin, Heidelberg: Springer Berlin Heidelberg), 493–495. doi: 10.1007/978-3-642-38954-2_137

Ruvindy, R., Barua, A., Bolch, C. J. S., Sarowar, C., Savela, H., and Murray, S. A. (2023). Genomic copy number variability at the genus, species and population levels impacts in situ ecological analyses of dinoflagellates and harmful algal blooms. ISME Communications 3, 70. doi: 10.1038/s43705-023-00274-0

Rysgaard, S., Mortensen, J., Juul-Pedersen, T., Sørensen, L. L., Lennert, K., Søgaard, D. H., et al. (2012). High air–sea CO2 uptake rates in nearshore and shelf areas of Southern Greenland: Temporal and spatial variability. Marine Chemistry 128–129, 26–33. doi: 10.1016/j.marchem.2011.11.002

Saiz, E., and Calbet, A. (2011). Copepod feeding in the ocean: scaling patterns, composition of their diet and the bias of estimates due to microzooplankton grazing during incubations. Hydrobiologia 666, 181–196. doi: 10.1007/s10750-010-0421-6

Saldarriaga, J. F., Taylor, F. J. R., Keeling, P. J., and Cavalier-Smith, T. (2001). Dinoflagellate Nuclear SSU rRNA Phylogeny Suggests Multiple Plastid Losses and Replacements. Journal of Molecular Evolution 53, 204–213. doi: 10.1007/s002390010210

Sassenhagen, I., Irion, S., Jardillier, L., Moreira, D., and Christaki, U. (2020). Protist Interactions and Community Structure During Early Autumn in the Kerguelen Region (Southern Ocean). Protist 171, 125709. doi: 10.1016/j.protis.2019.125709

Seelam, J. S., Fernandes De Souza, M., Chaerle, P., Willems, B., Michels, E., Vyverman, W., et al. (2022). Maximizing nutrient recycling from digestate for production of protein-rich microalgae for animal feed application. Chemosphere 290, 133180. doi: 10.1016/j.chemosphere.2021.133180

Seenivasan, R., Sausen, N., Medlin, L. K., and Melkonian, M. (2013). Picomonas judraskeda Gen. Et Sp. Nov.: The First Identified Member of the Picozoa Phylum Nov., a Widespread Group of Picoeukaryotes, Formerly Known as ‘Picobiliphytes.’ PLoS ONE 8, e59565. doi: 10.1371/journal.pone.0059565

Seuthe, L., Rokkan Iversen, K., and Narcy, F. (2011). Microbial processes in a high-latitude fjord (Kongsfjorden, Svalbard): II. Ciliates and dinoflagellates. Polar Biol 34, 751–766. doi: 10.1007/s00300-010-0930-9

Sherr, E. B., Sherr, B. F., Wheeler, P. A., and Thompson, K. (2003). Temporal and spatial variation in stocks of autotrophic and heterotrophic microbes in the upper water column of the central Arctic Ocean. Deep Sea Research Part I: Oceanographic Research Papers 50, 557–571. doi: 10.1016/S0967-0637(03)00031-1

Siano, R., Alves-de-Souza, C., Foulon, E., Bendif, E. M., Simon, N., Guillou, L., et al. (2011). Distribution and host diversity of Amoebophryidae parasites across oligotrophic waters of the Mediterranean Sea. Biogeosciences 8, 267–278. doi: 10.5194/bg-8-267-2011

Skovgaard, A. (2014). Dirty Tricks in the Plankton: Diversity and Role of Marine Parasitic Protists. Acta Protozoologica 2014, 51–62.

Smith, R. W., Bianchi, T. S., Allison, M., Savage, C., and Galy, V. (2015). High rates of organic carbon burial in fjord sediments globally. Nature Geosci 8, 450–453. doi: 10.1038/ngeo2421

Spring, S., Bunk, B., Spröer, C., Schumann, P., Rohde, M., Tindall, B. J., et al. (2016). Characterization of the first cultured representative of *Verrucomicrobia* subdivision 5 indicates the proposal of a novel phylum. The ISME Journal 10, 2801–2816. doi: 10.1038/ismej.2016.84

Stackebrandt, E. ed. (1991). *Nucleic acid techniques in bacterial systematics*. Chichester: Wiley.

Stoeck, T., Bass, D., Nebel, M., Christen, R., Jones, M. D. M., Breiner, H., et al. (2010). Multiple marker parallel tag environmental DNA sequencing reveals a highly complex eukaryotic community in marine anoxic water. Molecular Ecology 19, 21–31. doi: 10.1111/j.1365-294X.2009.04480.x

Stuart-Lee, A. E., Mortensen, J., Juul-Pedersen, T., Middelburg, J. J., Soetaert, K., Hopwood, M. J., et al. (2023). Influence of glacier type on bloom phenology in two Southwest Greenland fjords. Estuarine, Coastal and Shelf Science 284, 108271. doi: 10.1016/j.ecss.2023.108271

Stuart-Lee, A. E., Mortensen, J., Kaaden, A. -S. V. D., and Meire, L. (2021). Seasonal Hydrography of Ameralik: A Southwest Greenland Fjord Impacted by a Land-Terminating Glacier. JGR Oceans 126, e2021JC017552. doi: 10.1029/2021JC017552

Stuart-Lee, A., Møller, E. F., Winding, M., Van Oevelen, D., Hendry, K. R., and Meire, L. (2024). Contrasting copepod community composition in two Greenland fjords with different glacier types. Journal of Plankton Research 46, 619–632. doi: 10.1093/plankt/fbae060

Teeling, H., Fuchs, B. M., Becher, D., Klockow, C., Gardebrecht, A., Bennke, C. M., et al. (2012). Substrate-Controlled Succession of Marine Bacterioplankton Populations Induced by a Phytoplankton Bloom. Science 336, 608–611. doi: 10.1126/science.1218344

Teeling, H., Fuchs, B. M., Bennke, C. M., Krüger, K., Chafee, M., Kappelmann, L., et al. (2016). Recurring patterns in bacterioplankton dynamics during coastal spring algae blooms. eLife 5, e11888. doi: 10.7554/eLife.11888

Thiele, S., Vader, A., Thomson, S., Saubrekka, K., Petelenz, E., Armo, H. R., et al. (2023). The summer bacterial and archaeal community composition of the northern Barents Sea. Progress in Oceanography 215, 103054. doi: 10.1016/j.pocean.2023.103054

Thingstad, T. F., HagstrÖm, Å., and Rassoulzadegan, F. (1997). Accumulation of degradable DOC in surface waters: Is it caused by a malfunctioning microbialloop? Limnology & Oceanography 42, 398–404. doi: 10.4319/lo.1997.42.2.0398

Valencia, B., Stukel, M. R., Allen, A. E., McCrow, J. P., Rabines, A., and Landry, M. R. (2022). Microbial communities associated with sinking particles across an environmental gradient from coastal upwelling to the oligotrophic ocean. Deep Sea Research Part I: Oceanographic Research Papers 179, 103668. doi: 10.1016/j.dsr.2021.103668

Van Der Loos, L. M., D’hondt, S., Willems, A., and De Clerck, O. (2021). Characterizing algal microbiomes using long-read nanopore sequencing. Algal Research 59, 102456. doi: 10.1016/j.algal.2021.102456

Vaulot, D., Campo, J. D., Mahwash Jamy Burki, F., Guillou, L., Santoferrara, L., et al. (2023). pr2database/pr2database: PR2 version 5.0.0. doi: 10.5281/ZENODO.7805244

Von Scheibner, M., Sommer, U., and Jürgens, K. (2017). Tight Coupling of Glaciecola spp. and Diatoms during Cold-Water Phytoplankton Spring Blooms. Front. Microbiol. 8. doi: 10.3389/fmicb.2017.00027

Vonnahme, T. R., Chitkara, C., Krawczyk, D., Meire, L., Skogseth, R., Vader, A., et al. (2025). Abrupt decline of microplankton species richness linked to coastal inflow in an Arctic fjord. Limnology & Oceanography 70, 2688–2702. doi: 10.1002/lno.70159

Wadham, J. L., Hawkings, J. R., Tarasov, L., Gregoire, L. J., Spencer, R. G. M., Gutjahr, M., et al. (2019). Ice sheets matter for the global carbon cycle. Nat Commun 10, 3567. doi: 10.1038/s41467-019-11394-4

Wang, Z., Chang, X., Yang, X., Pan, L., and Dai, J. (2014). Draft Genome Sequence of Polaromonas glacialis Strain R3-9, a Psychrotolerant Bacterium Isolated from Arctic Glacial Foreland. Genome Announc 2, e00695–14. doi: 10.1128/genomeA.00695-14

Wei, T., and Simko, V. (2010). corrplot: Visualization of a Correlation Matrix. 0.95. doi: 10.32614/CRAN.package.corrplot

Wickham, H. (2016). ggplot2: elegant graphics for data analysis., Second edition. Cham: Springer international publishing.

Wilson, B., Müller, O., Nordmann, E.-L., Seuthe, L., Bratbak, G., and Øvreås, L. (2017). Changes in Marine Prokaryote Composition with Season and Depth Over an Arctic Polar Year. Front. Mar. Sci. 4. doi: 10.3389/fmars.2017.00095

Xue, C., Xie, Z.-X., Li, Y.-Y., Chen, X.-H., Sun, G., Lin, L., et al. (2021). Polysaccharide utilization by a marine heterotrophic bacterium from the SAR92 clade. FEMS Microbiology Ecology 97, fiab120. doi: 10.1093/femsec/fiab120

Yeh, Y.-C., and Fuhrman, J. A. (2022). Contrasting diversity patterns of prokaryotes and protists over time and depth at the San-Pedro Ocean Time series. ISME Communications 2, 36. doi: 10.1038/s43705-022-00121-8

Yih, W., and Coats, D. W. (2000). Infection of *Gymnodinium sanguineum* by the Dinoflagellate *Amoebophrya* sp.: Effect of Nutrient Environment on Parasite Generation Time, Reproduction, and Infectivity. J Eukaryotic Microbiology 47, 504–510. doi: 10.1111/j.1550-7408.2000.tb00082.x

Zapata, M., Fraga, S., Rodríguez, F., and Garrido, J. (2012). Pigment-based chloroplast types in dinoflagellates. Mar. Ecol. Prog. Ser. 465, 33–52. doi: 10.3354/meps09879

