## Supplementary materials for "Seasonal restructuring of heterotrophic microbial communities is differentially affected by glacier type in Greenland fjords"

**Table S1.** Sampling locations, including fjord, station name, geographic coordinates (latitude and longitude), sampling date, and depth (m) of the deep chlorophyll maximum (DCM).

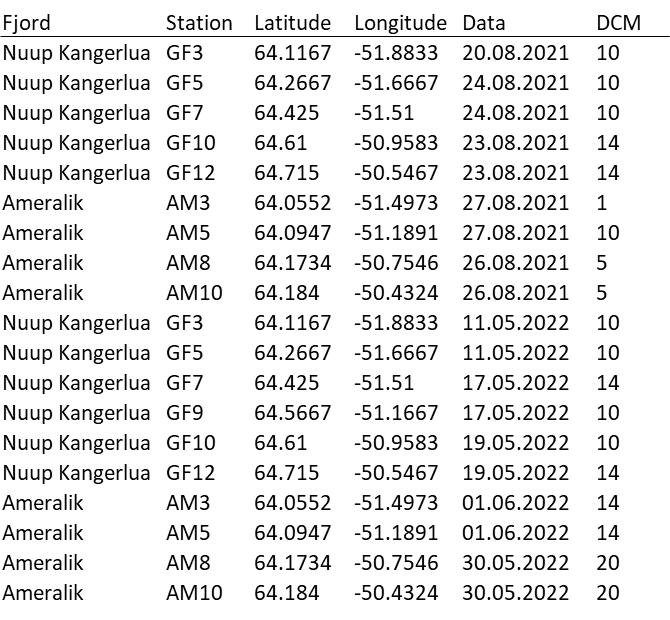

**S1 iFCM**

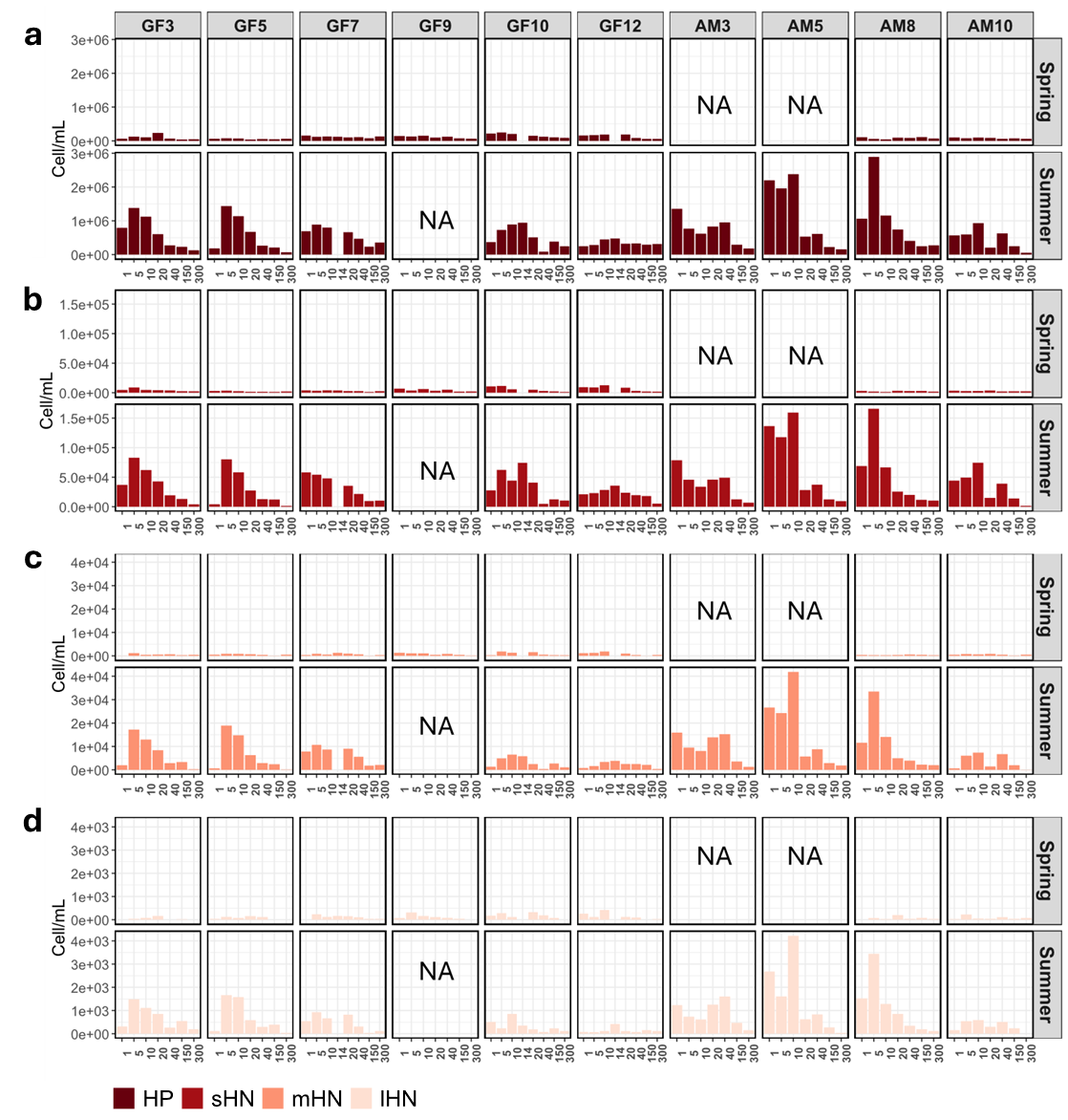

**Figure S1:** (a-b-c-d) Bar plots of microbial abundances (cells mL-1) at discrete sampling depths in NK (GFx) and AM (AMx) in spring (top) and summer (bottom). (a) HP – bacteria/heterotrophic picoplankton – dark red, (b) sHN – small heterotrophic nanoplankton – brick red, mHN – medium heterotrophic nanoplankton– light orange, lHN – large heterotrophic nanoplankton – pale peach. DCM = deep chlorophyll maximum. Note different scale of y-axis. NA – data not available.

In spring, heterotrophic picoplankton (HP) was most abundant in NK, particularly in the inner fjord (Fig. S1a). HP concentrations averaged ~1.1 × 10⁵ cells mL⁻¹ in NK and ~8.0 × 10⁴ cells mL⁻¹ in inner AM, peaking at ~2.5 × 10⁵ cells mL⁻¹ in surface waters (1–10 m) at GF10. Small, medium, and large heterotrophic nanoplankton (sHN, mHN, and lHN, respectively) were most abundant in surface waters (1–20 m) at GF10 and GF12, averaging ~9.0 × 10³, ~1.0 × 10³, and ~2.0 × 10² cells mL⁻¹, respectively (Fig. S1b-d). Cell concentrations in the outer fjord were approximately half of those observed in the inner fjord and were comparable to those in inner AM (no data for outer AM). In summer, heterotrophic groups increased up to threefold in AM5 and AM8 compared to NK, where the lowest concentrations were observed at GF12 (Fig. S1a-d). In NK, HP averaged ~5.0 × 10⁵ cells mL⁻¹, whereas in AM8 its abundance peaked at ~3.0 × 10⁶ cells mL⁻¹ at 5 m depth, with an overall fjord-wide average of ~8.0 × 10⁵ cells mL⁻¹. Both sHN and lHN increased by approximately one order of magnitude in summer compared to spring, while mHN concentrations remained similar in NK but were higher in surface waters (1–10 m) at AM5 and AM8.

**S2 Metabarcoding**

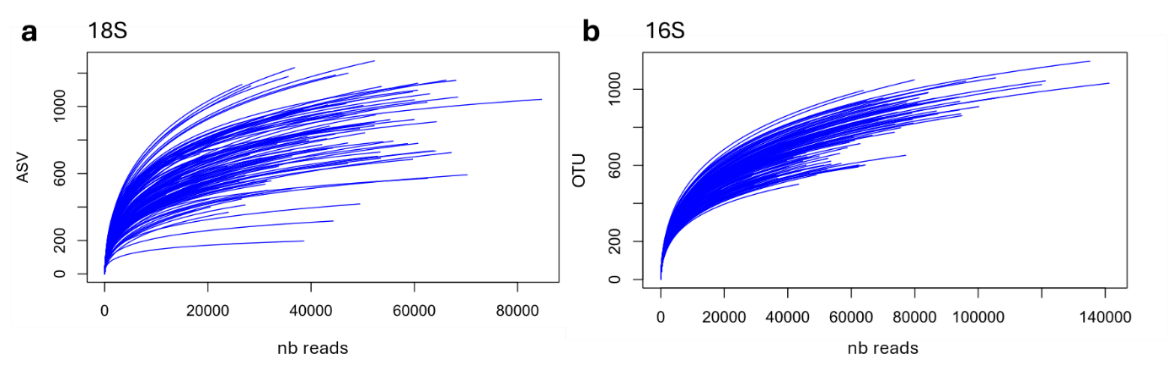

**Figure S2:** Rarefaction curves of (a) V4 region 18S rRNA – heterotrophic protist and (b) full 16S rRNA gene.

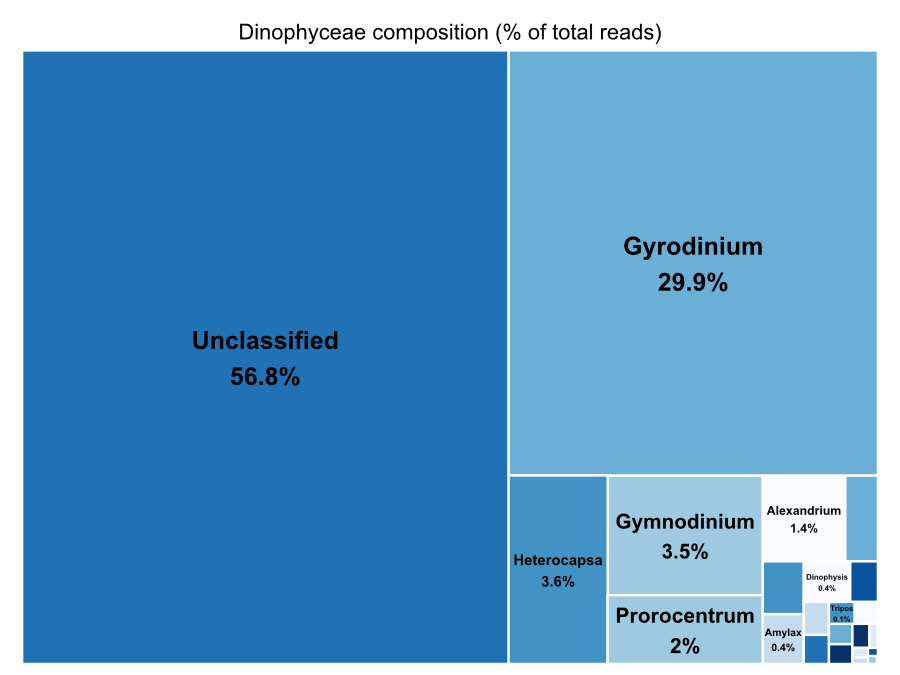

**Figure S3.** Treemap showing the relative abundance (%) of Dinophyceae genera across all samples. The area of each rectangle is proportional to the total number of sequences assigned to each genus. Percentages indicate the contribution of each genus to the total Dinophyceae reads.

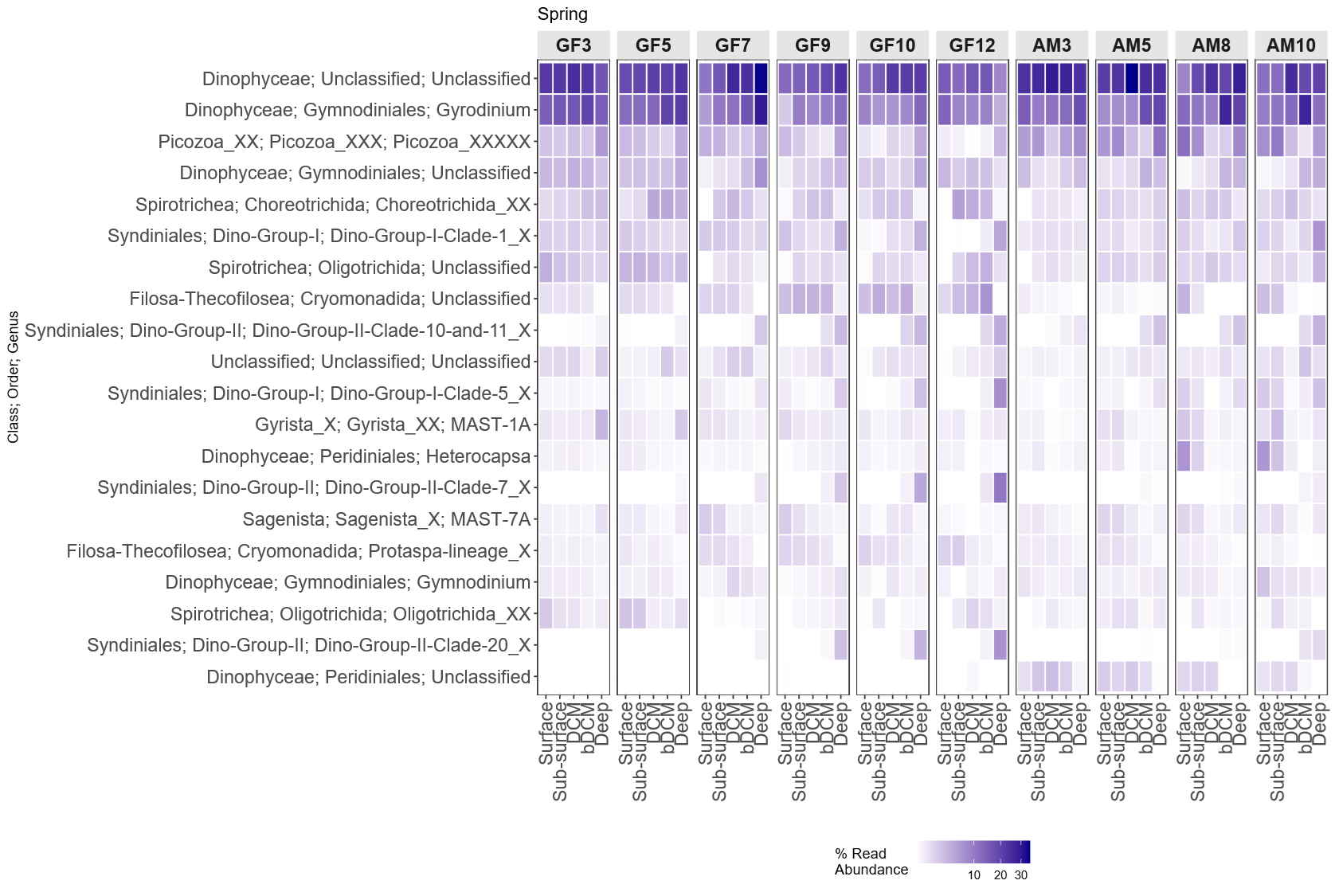

**Figure S4:** Heatmap of the relative abundance of the 20 most abundant heterotrophic protist taxa detected in spring. Taxa are labeled by class, order and genus. X-axis labels indicate sampling depths: Surface = 1 m; Subsurface = 5 and 10 m; DCM = deep chlorophyll maximum; bDCM = 20 and 40 m; Deep = 150 and 300 m. Station codes: GFx = NK fjord; AMx = AM fjord.

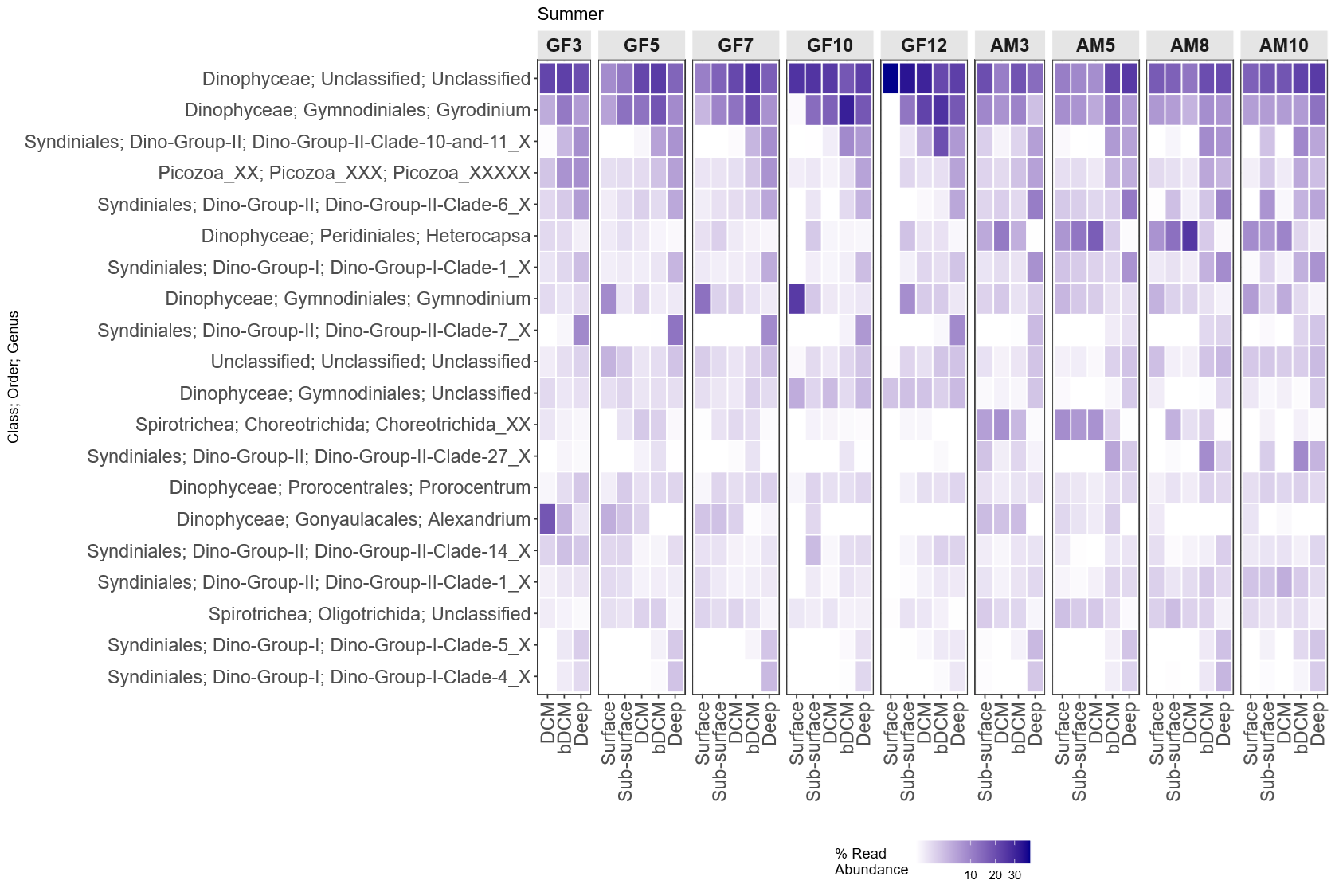

**Figure S5:** Heatmap of the relative abundance of the 20 most abundant heterotrophic protist taxa detected in summer. Taxa are labeled by class, order and genus. X-axis labels indicate sampling depths: Surface = 1 m; Subsurface = 5 and 10 m; DCM = deep chlorophyll maximum; bDCM = 20 and 40 m; Deep = 150 and 300 m. Station codes: GFx = NK fjord; AMx = AM fjord.

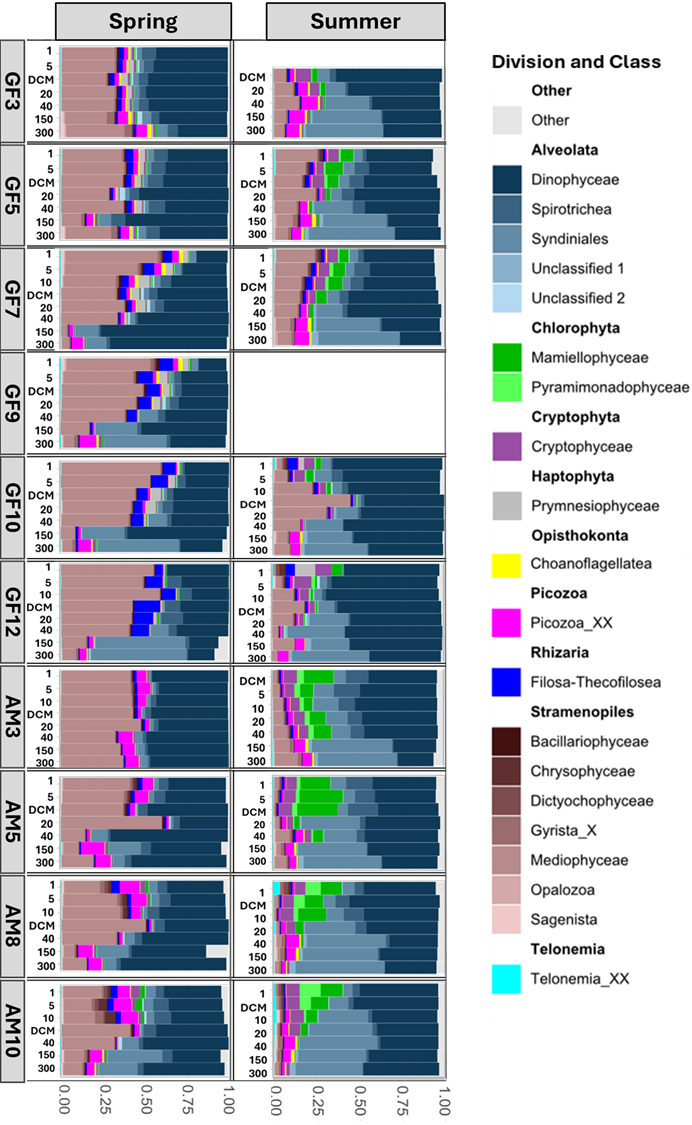

**Figure S6:** Nested bar plots of protist relative abundances by division and 20 most abundant classes (~96% of total reads) across discrete depths and stations in NK (GFx) and AM (AMx) during spring (left) and summer (right), progressing from outer (GF3, AM3) to inner fjord (GF12, AM10). Other- represent less abundant classes.

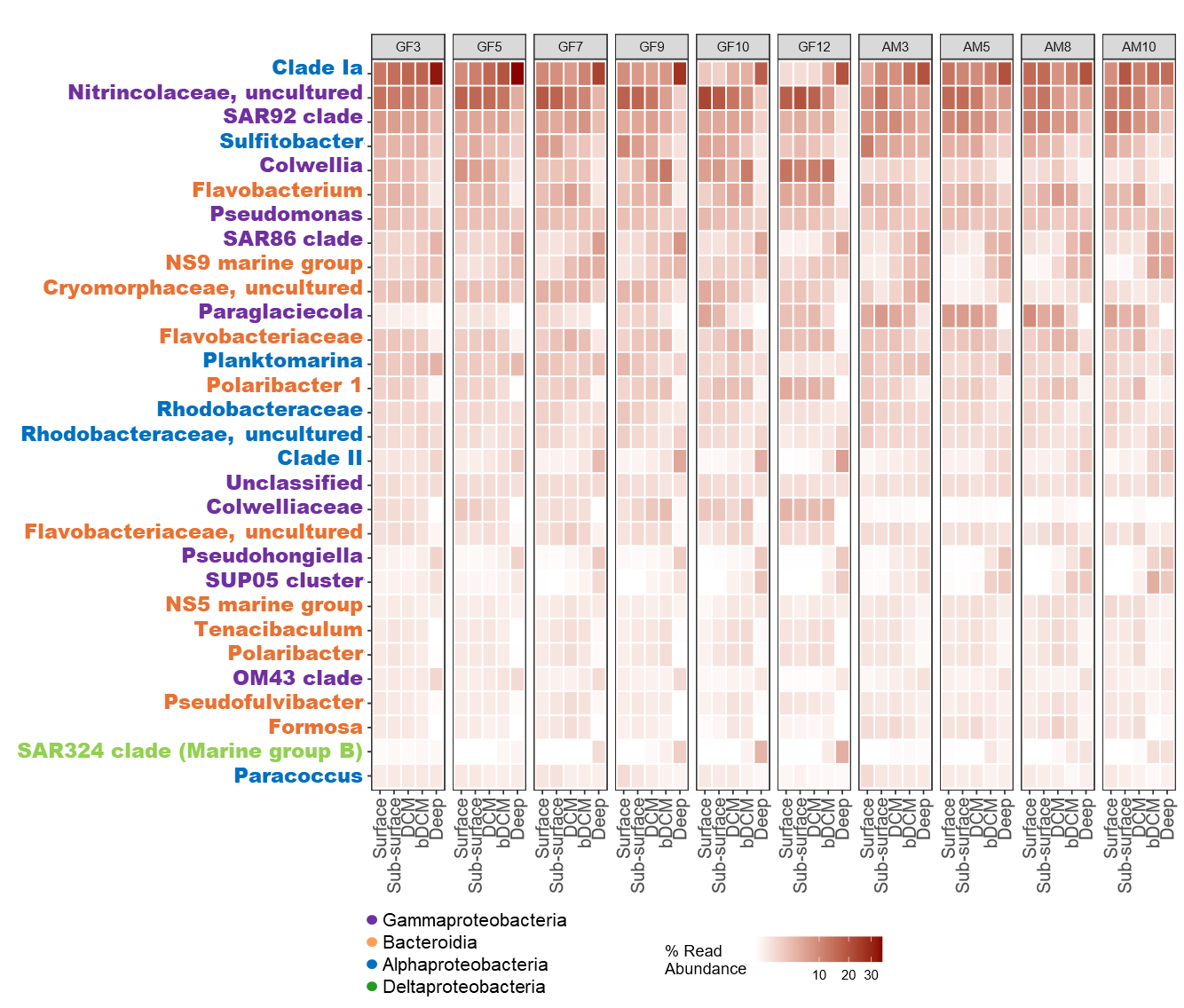

**Figure S7:** Heatmap of the relative abundance of the 30 most abundant bacterial taxa detected in spring. Taxa are labeled by genus or the lowest taxonomic level identified and colored by Class. X-axis labels indicate sampling depths: Surface = 1 m; Subsurface = 5 and 10 m; DCM = deep chlorophyll maximum; bDCM = 20 and 40 m; Deep = 150 and 300 m. Station codes: GFx = NK fjord; AMx = AM fjord.

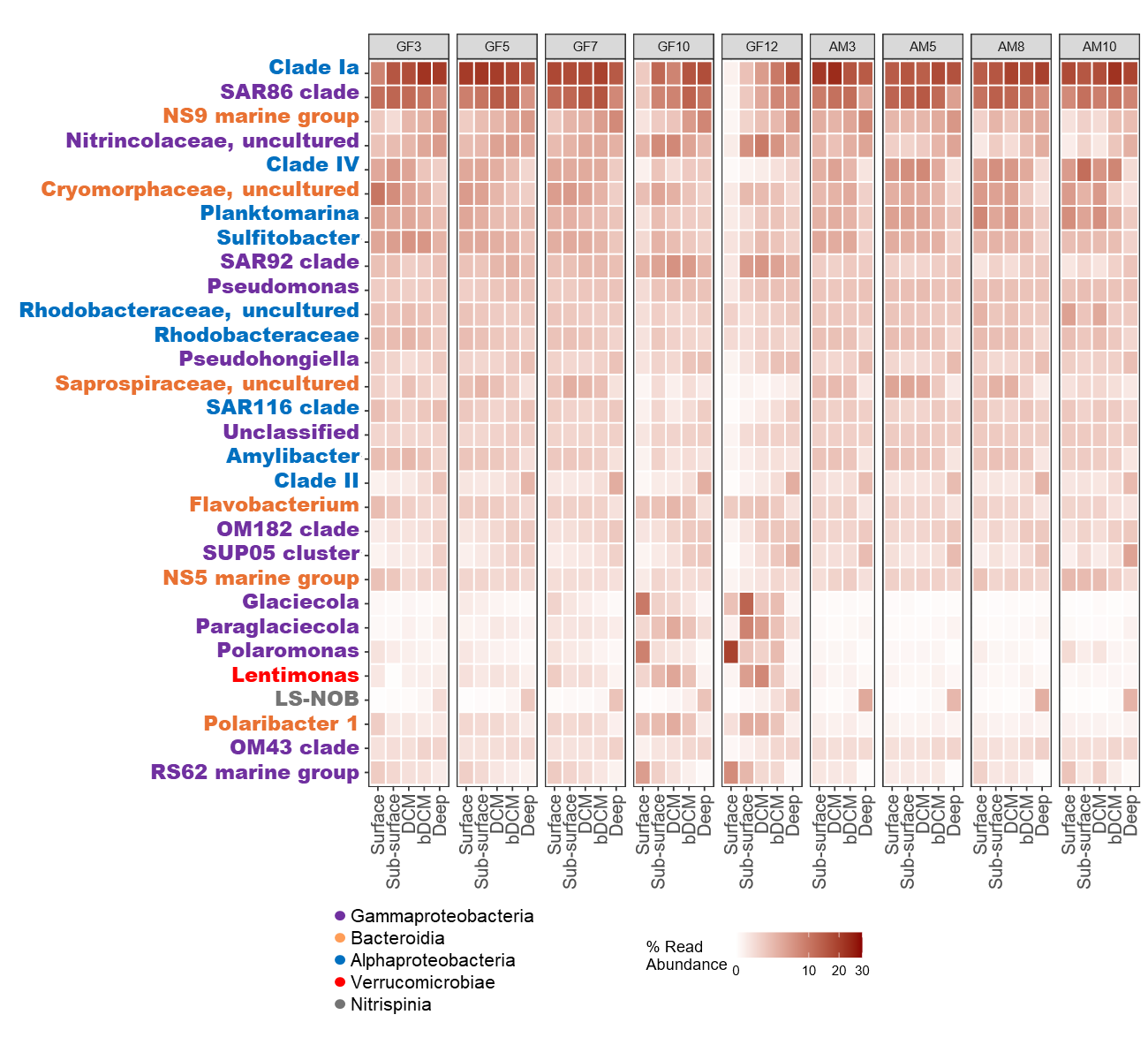

**Figure S8:** Heatmap of the relative abundance of the 30 most abundant bacterial taxa detected in summer. Taxa are labeled by genus or the lowest taxonomic level identified and colored by Class. X-axis labels indicate sampling depths: Surface = 1 m; Subsurface = 5 and 10 m; DCM = deep chlorophyll maximum; bDCM = 20 and 40 m; Deep = 150 and 300 m. Station codes: GFx = NK fjord; AMx = AM fjord.

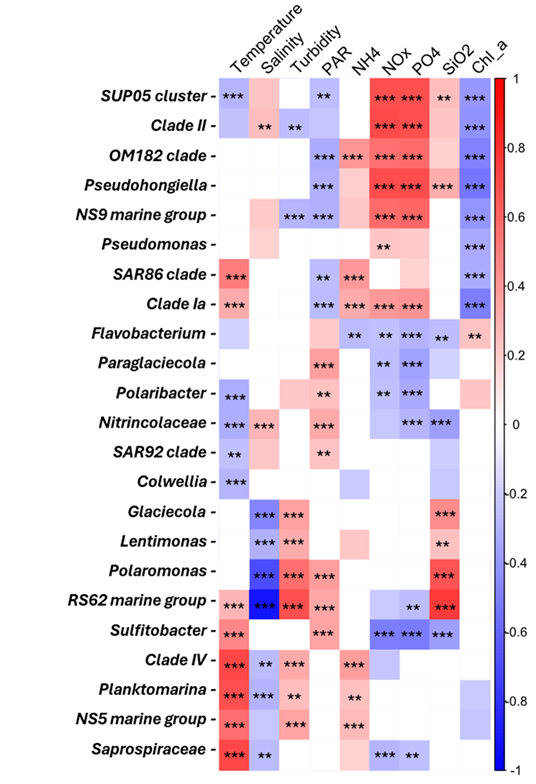

**Figure S9.** Corrplot showing Pearson’s correlation matrix between the most abundant taxa and environmental variables. Red indicates positive correlations, blue negative correlations. Only significant corr elations are displayed (p < 0.05). Significance levels: *** p < 0.001; ** p < 0.005.
